# Stress triggers global histone synthesis in the liver

**DOI:** 10.64898/2026.09.22.753600

**Authors:** Shreenidhi Rajkumar, Lavanya Vumma, Jacinth Naidoo, Min Zhu, Ishrat Durdana, Sri Vidhya Chandrasekar, Caiden Golder, Harsh Goar, W. Kiran Kicinski, Rico Gamuyao, Yudong Wei, Hao Zhu, Suraj J. Patel, Joseph M. Ready, Michael Buszczak, Joshua J. Gruber

## Abstract

Despite extensive drug development targeting DNA synthesis, there have been few efforts to inhibit histone synthesis, which occurs jointly with DNA synthesis in S-phase. A major roadblock is the lack of tools to quantitively measure histone synthesis due to the high abundance and stability of histone proteins. Here we present chemical labeling methodologies to produce the first *in vivo* measurements of histone synthesis rates. Under basal conditions, histone synthesis tracks closely with tissue proliferation. However, multiple systemic and liver-specific stresses induce a previously uncharacterized surge of hepatocyte nucleosome production that is uncoupled from DNA replication. Tracing of newly synthesized nucleosomes detected promoter deposition, which was required to buffer transcriptional output. These results may inform strategies to modulate histone production to influence stress responses or disease.

## Main Text

Multiple highly active cancer therapeutics target S-phase processes including nucleotide synthesis, nucleotide salvage, DNA unwinding, replication and others. Histone proteins are the major protein class synthesized to generate nucleosomes to support DNA replication in S-phase. Despite the strong rationale to target histone synthesis as a potential cancer therapeutic strategy there remains a lack of chemical and genetic strategies to validate this concept. A major roadblock is the difficulty of measuring histone synthesis in cells and animals. This is because core histones (H3, H4, H2A, H2B) are highly abundant nuclear proteins, comprising ~1% of the dry weight of mammalian cells, and have one of the longest protein half-lives, with estimates exceeding one year in non-dividing cells (*1–3*). Their abundance and stability create limitations for accurately measuring their levels, synthesis (*4*), and degradation (*5*). Given the critical role of histones in DNA compaction, gene regulation and genome stability, increased efforts to understand how histone synthesis and stability is regulated should have widespread importance and would also inform strategies to target their production for therapeutic value.

Our recent work identified that histone chaperones that guide the maturation and deposition of histone H3/H4 dimers are critical for maintaining histone supply in replicating cells (*6*). This has focused attention on whether targeting histone supply could be an exploitable dependency in human cancers or other diseases. However, methods to measure and quantify histone production are limited, which constrains the ability to design inhibitors to this pathway (*7*). Recent work using chemical labeling tags to trace histone incorporation during DNA replication in cell lines (*8*) utilized transgenes driven by heterologous promoters that may not accurately reflect endogenous regulation. Similarly, common genetic lineage tracing methods like *cre-loxp* systems (*9*) which can be induced *in vivo* lack the simplicity, time-resolution and versatility to measure protein dynamics across a wide variety of tissues in unison. Also, previous chemical labeling strategies lack specificity, or induce toxicity, which limits widespread implementation (*10–14*). Therefore, real-time, quantitative protein labeling *in vivo* has thus far yet to be demonstrated, a key gap that is addressed by the current work.

Herein, we introduce chemical tools, methodologies, and genetic strategies, which when combined, allow for robust, time-resolved endogenous protein labeling and quantitation *in vivo*. We apply these methods to directly measure histone synthesis rates *in vivo* for the first time. We make the unexpected discovery of non-replication-dependent production of core histones in response to liver stress and use our methods to dissect the regulatory principles and chromatin impacts of this phenomenon. Taken together, these works outline general principles for measuring protein production rates *in vivo* which could be adapted to any protein-of-interest.

### Characterization of *SNAP-H4C3* mouse model and SNAP-labeling substrates

SNAP is an engineered protein domain derived from human O6-benzylguanine DNA alkyl-transferase that forms a self-labeling covalent adduct to benzylguanine-containing analogs (*15*) (**Fig. S1a**). When appended to proteins, SNAP tags offer several features ideal for *in-vivo* applications, including rapid and efficient one-step labeling, minimal off-target reactivity, low cellular toxicity. A wide array of cell-permeable SNAP-tag substrates are commercially available (*16*) and customizable. To ensure proper endogenous regulation we used CRISPR/Cas9 to precisely insert a SNAP tag in-frame at the N-terminus of a single allele of the mouse *H4c3* gene, which encodes a prototypical and widely expressed replication-dependent H4 gene in the major histone locus (Hist1 locus) on chromosome 13 (**Fig. 1a**). We detected robust expression of the SNAP-H4 protein across multiple tissues (**Fig. S1b**). Body weights measured at 10 weeks of age showed no significant differences between heterozygous (*Snap-H4c3*) and wild-type (*WT*) littermates (**Fig. S1c**). Colony breeding followed expected Mendelian ratios (**Fig. S1d**). To confirm chromatin-incorporation of SNAP-H4, we acid-extracted histones from mammalian embryonic fibroblasts (MEFs) followed by immunoblotting with an anti-SNAP antibody. SNAP-H4 was detected in the chromatin fraction (**Fig. S1e**), indicating proper maturation and processing of the tagged histone.

**Fig. 1:**
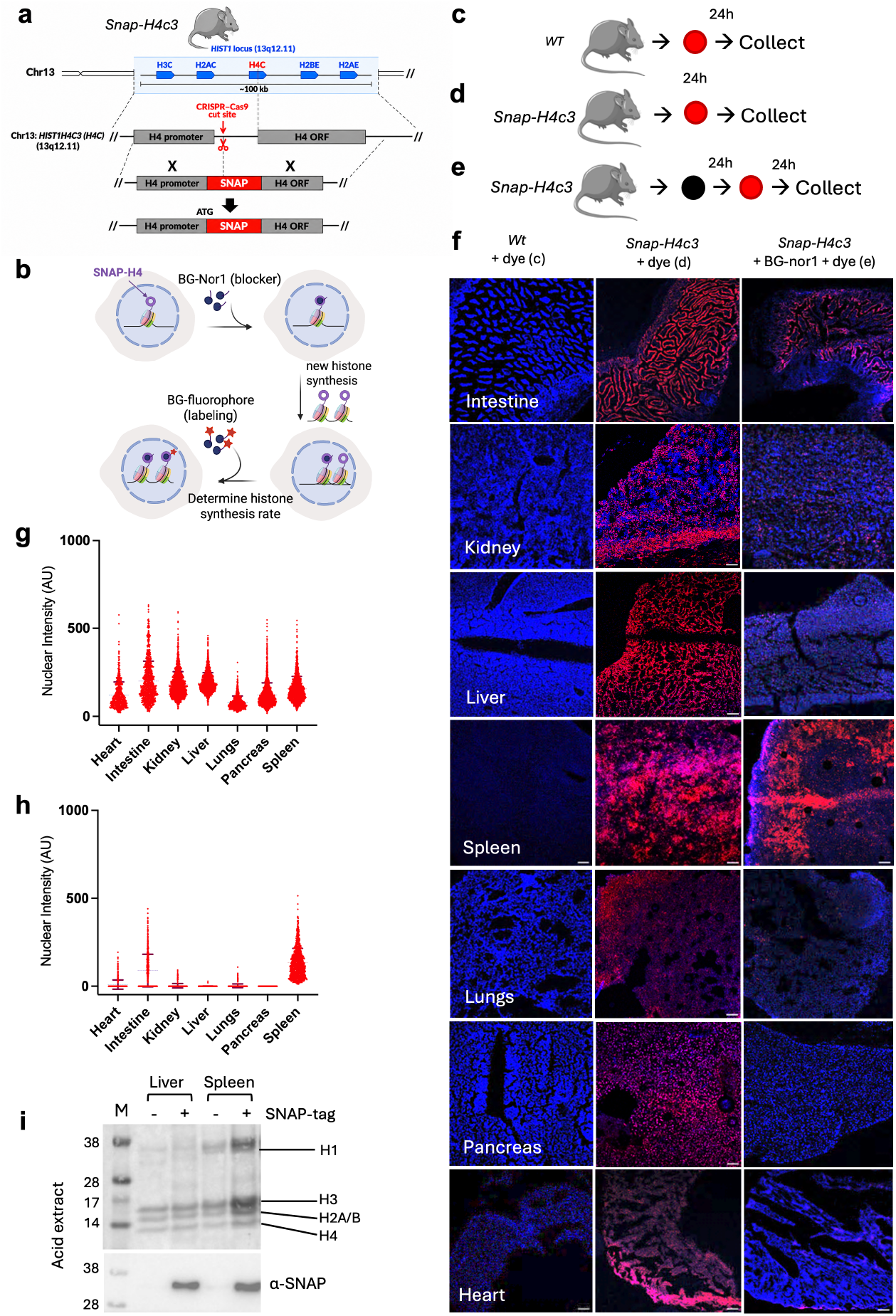
*In vivo* SNAP labeling of major tissues to determine histone synthesis rates. **a**, Schematic showing SNAP-covalent labeling. **b**, CRISPR–Cas9 targeting strategy for SNAP insertion at the *h4c3* locus, chr13. **c, d, e** Schematic of *in vivo* labeling strategy to detect histone H4 in *WT* vs *Snap-H4c3* animals (n = 2) and to detect newly synthesized histone H4 in 48h using pulse-chase strategy. TMR-Star (red),4B0G-Nor1 (black) **f**, Results of SNAP labeling *in vivo* across various tissues. Tissues were harvested, flash frozen, cryo-sectioned, stained with DAPI & imaged at 570 nm by confocal; scale bar 10 μm, imaged at 10x magnification. Each column corresponds to different labeling strategy. **g**, Quantification of results from (d); nuclear intensity per nuclei was quantified, plotted as column individual scatter plot (median); n = 2 per column. **h**, Quantification of results from strategy (e); nuclear intensity per n4u5clei was quantified, plotted as column individual scatter plot (median); n = 2 per column. **i**, Acid-extracted histones from *WT* and *Snap-H4c3* tissues. Total histone levels stained with Memcode blue, then the membrane was destained and probed with anti-SNAP antibody

For *in vivo* labeling, we used TMR-Star as the fluorescent SNAP substrate (**Fig. S1f**) and BG-Nor1 (*17*)as a non-fluorescent blocking reagent (**Fig. S1g**). TMR-Star exhibits an excitation maximum at 554 nm and an emission maximum at 580 nm, enabling robust fluorescent detection. BG-Nor1, by contrast, is non-fluorescent and serves to irreversibly occupy pre-existing SNAP-tagged histones. This combination allows for a pulse-chase strategy (*5*), in which pre-existing (“old”) histones are blocked with BG-Nor1, followed by a chase period, after which newly synthesized histones can be selectively labeled with TMR-Star (**Fig. 1b)**. Next, we evaluated the pharmacokinetic properties of TMR-Star and BG-Nor1 to guide *in vivo* dosing parameters. TMR-Star exhibited approximately twice the half-life (T1/2) and peak plasma concentration (C_max_) compared to BG-Nor1 (**Fig. S1h**,**i**). Based on these data, we selected *in vivo* dosing regimens of 5 mg/kg i.p. for TMR-Star and 10 mg/kg i.p. for BG-Nor1. Both compounds were effectively cleared from circulation by 24 hours post-injection (**Fig. S1h**), thereby establishing a 24-hour period for both blocking and labeling.

To validate the performance of the BG-Nor1 blocker, we derived and immortalized MEFs from littermate *WT* and *Snap-H4c3* embryos. Immortalized *Snap-H4c3* MEFs were labeled with TMR-Star *in vitro* to assess fluorescent substrate efficiency. To evaluate blocking efficacy, cells were pre-treated with either in-house synthesized BG-Nor1 or a commercially available NEB blocking reagent prior to TMR-Star labeling. TMR-Star showed robust labeling performance *in vitro*, while BG-Nor1 was equally as effective as NEB commercial blocker in occupying the tags (**Fig. S1j**,**k**), confirming its suitability for further *in vivo* studies.

To evaluate genome-wide incorporation of the SNAP-H4 histone, we performed nucleosome-seq after MNase digestion from SNAP-H4-continaing MEFs. We sequenced all input nucleosomes, as well as nucleosomes specifically containing SNAP-H4 by capturing them on BG-beads followed by vigorous washing. Extensive overlap of the dyad density maps between input & SNAP-pulldown (**Fig. S1l**) confirmed that SNAP-H4 incorporation does not disturb nucleosome occupancy or positioning. Similarly, we observed scant differences in gene expression between *WT* and *Snap-H4c3* mouse livers (**Fig. S1m**). Taken together, the SNAP-H4 tracer appeared to follow typical chromatin integration patterns without perturbing chromatin architecture or gene regulation, making it a suitable system to analyze histone production rates.

### Measuring global histone synthesis rates *in vivo*

First, we determined SNAP-H4 abundance in all tissues by administering a single intraperitoneal (i.p.) dose of TMR-Star to *WT* control (**Fig. 1c**) and *Snap-H4c3* mice (**Fig. 1d**). After 24 hours, all major organs were harvested and cryosectioned. Widespread labeling of histone SNAP-H4 was detected across all major tissues in *Snap-H4c3* mice, but not *WT* controls (**Fig. 1f**), confirming dye specificity and reflecting broad expression from the endogenous *H4c3* locus (**Fig. 1g**). A single dose of TMR-Star was sufficient to fully label all endogenous SNAP proteins, as experiments testing additional doses did not further boost signal (**Fig. S2a**,**b**,**c**). SNAP labeling did not perturb tissue architecture (**Fig. S2d**). Thus, real-time labeling of endogenous protein was robust, safe, and easily saturated.

Next, we applied a pulse-chase strategy to selectively label newly synthesized histones (**Fig. 1e**). Mice were first injected with BG-Nor1 to block existing (old) SNAP-H4. After 24 hours, TMR-Star was administered to label newly synthesized histones, and tissues were harvested 24 hour later. We observed strong TMR signal in the spleen and modest signal in the intestine (**Fig. 1f**), while most other organs showed little to no detectable labeling (**Fig. 1h**). In the intestine, labeling is likely due to ongoing epithelial proliferation, while in the spleen, labeling was concentrated in follicular zones, consistent with active proliferation of immune cell populations and their high demand for histone synthesis to support DNA replication (*18,19*). To confirm that the SNAP-H4 tracer was deposited into chromatin, histones were acid-extracted from liver and spleen and immunoblotted with an anti-SNAP antibody (**Fig. 1i**), revealing robust SNAP signal in nucleosomal histones.

To assess the long-term residence of labeled histones, we administered a single dose of TMR-Star then collected tissue samples 1- and 2-weeks post-injection (**Fig. S3a**). The fluorescence signal had largely dissipated in most tissues by 1 and 2 weeks, suggesting either active histone turnover or metabolism of the dye (**Fig. S3c**). In contrast, the spleen retained distinct clusters of brightly labeled cells, indicating that dye metabolism or histone turnover may be minimal in certain tissues.

The residence of the BG-Nor1 blocker was also assessed in an inverse experiment. Mice were first administered BG-Nor1 to irreversibly block pre-existing SNAP-tagged histones, followed by a 1-week and a 2-week waiting period. Subsequently, TMR-Star was administered 24 hours prior to tissue collection to label only newly synthesized histones (**Fig. S3b**). TMR signal was brighter in most tissues after 2 weeks versus 1 week (**Fig. S3c**), indicating that BG-Nor1 continued to block SNAP-H4 sites for at least one week *in vivo*.

### *Snap-H4c3* Reporter Reveals Non-Canonical Histone Production

We sought to extend the *Snap-H4c3* model by measuring histone production in cells induced to proliferate from a quiescent state *in vivo*. Hepatocytes are largely quiescent but can be induced to enter the cell cycle after liver injury to regenerate the organ. For example, after carbon tetrachloride (CCl4)-induced acute liver injury approximately 20–30% of hepatocytes are induced to proliferate at 48 hours (*20*) (**Fig. 2a**). To determine if these proliferating cells were also synthesizing new histones, we first blocked all pre-existing histones in *Snap-H4c3* animals, then administered CCl4 followed by labeling with TMR-Star and harvested after 24 or 48 hours (**Fig. 2b**). Unexpectedly, we observed that >90% of hepatocytes were labeled by TMR-Star after CCl4 treatment (**Fig. 2c,e**), consistent with widespread histone production and far outstripping the 20-30% expected to proliferate to enable regeneration. This suggested hepatocytes may produce histones in response to injury in a manner that could be uncoupled from cell division.

**Fig. 2:**
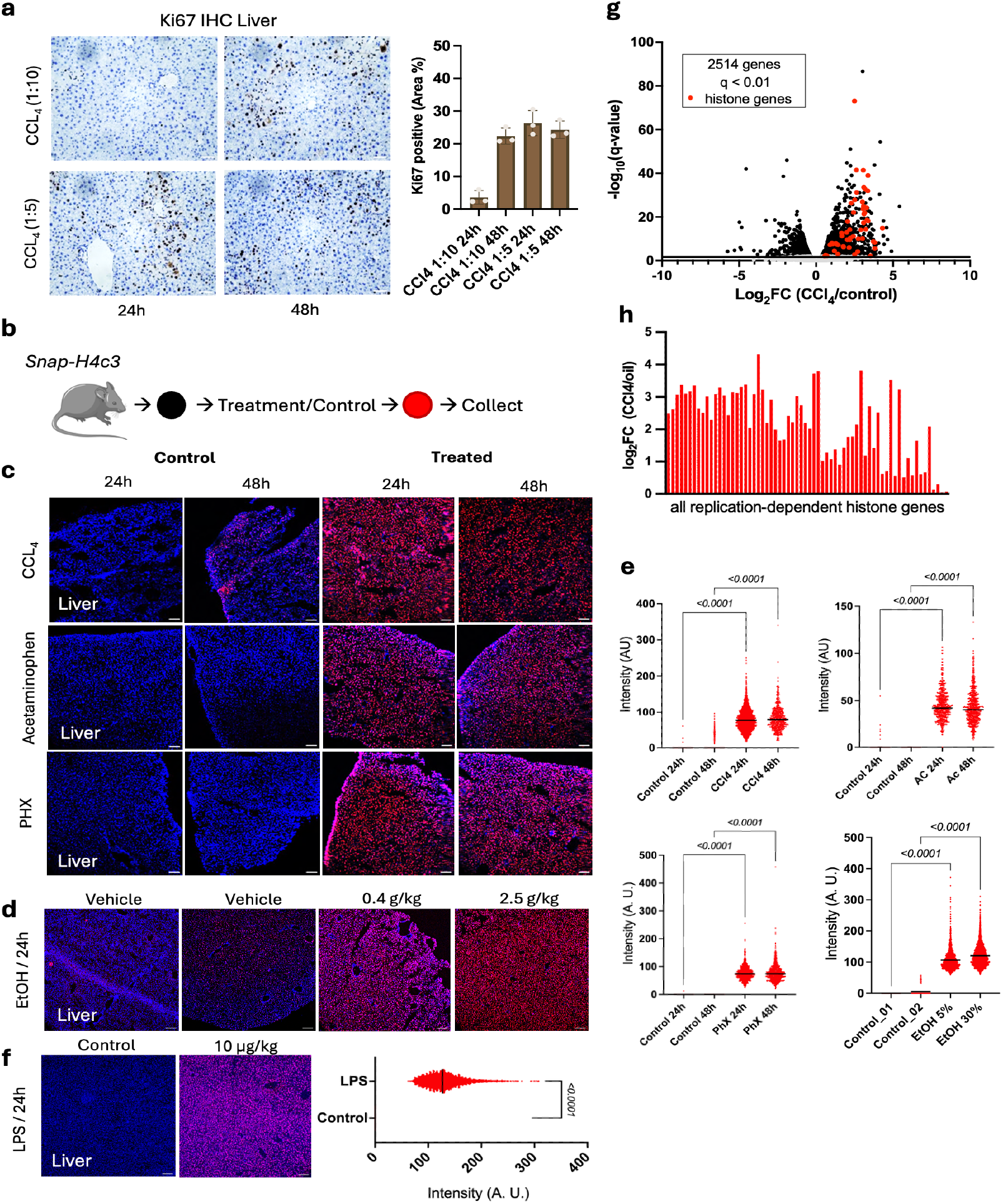
*Snap-H4c3* reporter reveals surge in stress–driven histone production. **a**, Ki67 IHC staining of CCl4 treated liver for 24 & 48h with quantification. 1:10 and 1:5 are CCl4 dilutions. **b**, Schematic for liver treatments: old histones were blocked by BG-Nor1 (black), chemical or physical damage is induced, and then TMR-star (red) labels all newly made histones. **c**, Results of SNAP labeling *in vivo* from different liver damaging agents (n = 2 for each condition). Tissues were harvested, flash frozen, cryo-sectioned, stained with DAPI & imaged at 570 nm by confocal; scale bar 10 μm, imaged at 10x magnification. **d**, Ethanol challenge at 0.4 g/kg (5%, x4 every 6h) and 2.5 g/kg (30%, once), all for 24h. Same experimental design and procedure followed as (c). **e**, Quantifications for (c & d) where each graph corresponds to each row of images. Nuclear intensity per nuclei was quantified, plotted as column individual scatter plot (median); n = 2 per column, two-way ANOVA (column-wise comparisons). **f**, LPS challenge at 10 ug/kg (i.p. once), for 24h. Same experimental design and procedure followed as (b, c). Nuclear intensity per nuclei was quantified, median indicated; n = 2 mice, welch’s test. **g**, rRNA depletion RNA-seq volcano plot shows histone gene induction (red). **h**, all replication-dependent histone transcripts are upregulated in comparison to oil controls.

To determine if this surge in histone production was specific to CCl4-mediated liver injury, or reflective of a generalized response, we tested other models of acute liver injury. After acetaminophen (Ac) treatment and partial hepatectomy (PHX) (**Fig. 2c,e**) we also observed that over 90% of hepatocytes produced new histones. Also, Ki67 immuno-histochemistry (IHC) of liver sections from mice treated with Ac and PHX (**Fig. S4a**,**b**) further confirmed that <20% of cells were proliferative, which also did not coincide with the widespread histone synthesis observed. Thus, extensive histone production in liver was observed after multiple types of liver insult and did not correlate with markers of proliferation.

To further distinguish the effects of stress-induced histone production from proliferation associated with regeneration we tested acute ethanol administration, which is known to impair the hepatocyte proliferation (*21*). Ethanol administration of 0.4 g/kg and 2.5 g/kg strongly stimulated global hepatic histone production in mice (**Fig. 2d,e**). Similarly, humans with alcohol-related steatosis (AS), alcoholic liver disease (ALD) with fibrosis (ALD-fib), and ALD with alcoholic hepatitis (AH) had elevated histone mRNA transcripts compared to healthy controls (**Fig. S4c,d**). To further test whether stress-induced histone production extends to non-proliferative inflammatory stimuli, we administered lipopolysaccharide (LPS), a potent inducer of acute systemic inflammation (*22*). LPS strongly stimulated hepatic histone production (**Fig. 2f**). This suggested the existence of a generalized stress-induced histone synthesis response in the liver that may be uncoupled from hepatocyte proliferation.

Next, we sought to confirm that the labeling of SNAP-H4 was a marker of global synthesis of core histones in response to liver stress, and not only expression from the *Snap-H4c3* reporter locus. Transcriptome analysis of rRNA-depleted total RNA showed that nearly 20% of all significantly upregulated transcripts corresponded to core replication-dependent histone genes (**Fig. 2g**). When the replication-dependent histone transcripts were specifically queried, we observed they were universally upregulated by CCl4 treatment (**Fig. 2h**). At the protein level, pulse-labeling with a click chemistry-enabled methionine analog (HPG) followed by biotin-coupling and streptavidin enrichment, revealed induction of newly synthesized histone proteins after CCl4 treatment in hepatocytes (**Fig. S4e**), thus confirming increased global histone production with an orthogonal technique. Together, these data validate that hepatocytes induce a program of global core histone synthesis after stress.

Although global histone synthesis appeared to be induced in liver after various stresses, there was a disconnect between the near-universal histone production, which is classically linked to S-phase progression, and the low levels of hepatocyte proliferation observed. To more directly test whether DNA replication accompanies hepatocyte histone production, we first assayed EdU incorporation after CCl4 treatment in WT animals. We found that only ~10% of hepatocytes showed EdU incorporation (**Fig. 3a**). Next, EdU and SNAP labeling were performed in unison following CCl4 treatment and scored in the same liver, intestine and spleen tissues (**Fig. 3b**). Serial EdU injections every 6 hours were performed to ensure labeling of all S-phase cells over the 24-hour period. In spleen and intestine, the overlap of SNAP-TMR signal with EdU was substantial, indicating tight correlation between DNA replication and histone synthesis (**Fig. 3c,d**). However, in CCl4-treated liver the proportion of cells undergoing DNA replication and histone synthesis jointly was much more limited (**Fig. 3c,d,e**), as most cells producing histones did not take up EdU. Finally, we did not detect a correlation between hepatocyte DNA damage, as measured by γ-H2AX staining, and histone production after injury (**Fig. S4f,g**). We conclude that after acute stress the histone synthesis response that occurs in hepatocytes is uncoupled from DNA synthesis or DNA repair.

**Fig. 3:**
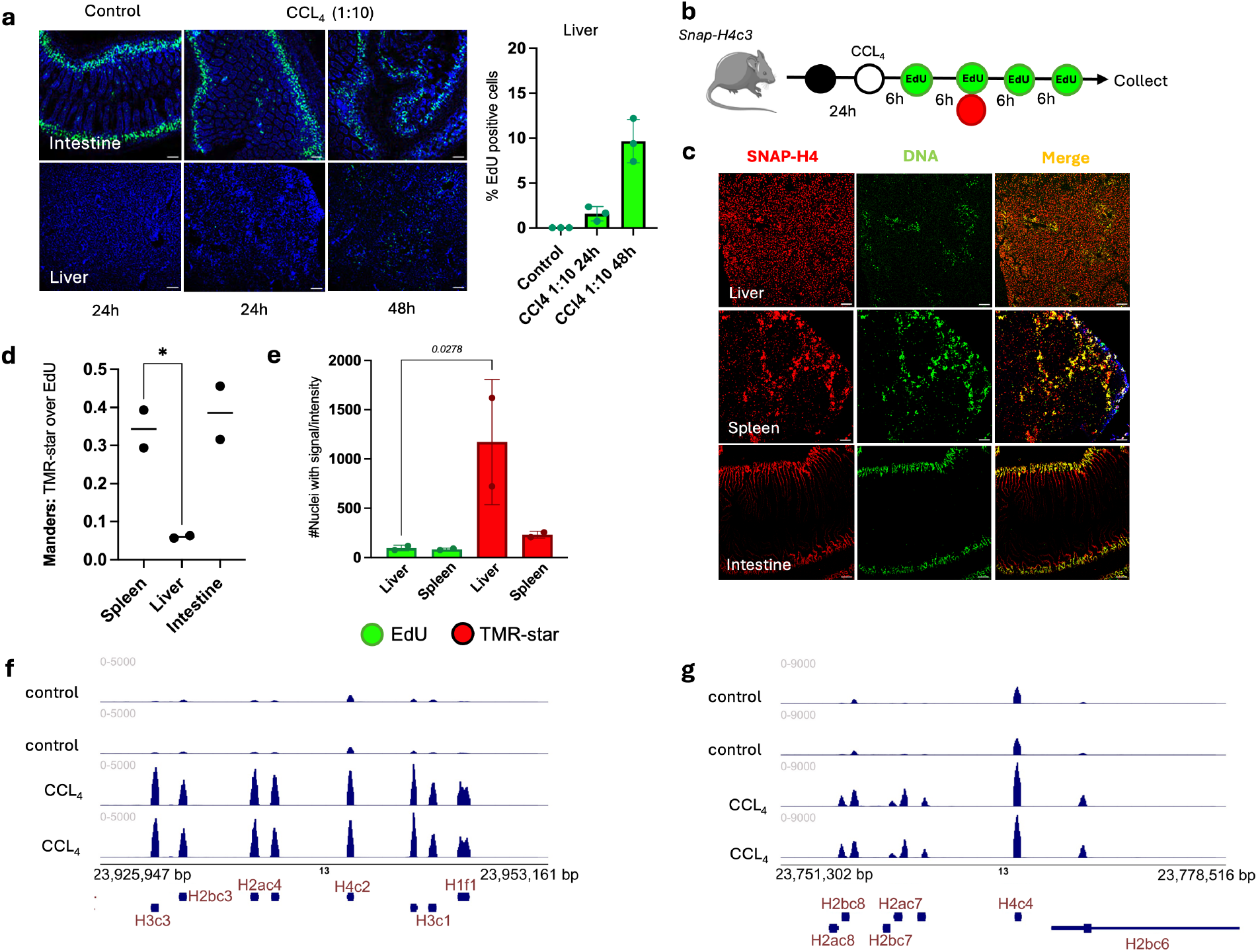
Histone synthesis is uncoupled from DNA replication after liver injury. **a**, EdU assay (Alexa488) on intestine & liver from *WT* animals treated with CCl4 (1:10 dilution in vehicle, n = 2), scale bar 10 μm, imaged at 10x magnification and quantification: EdU staining was quantified as the percentage of positively stained area relative to the total tissue area (area %). **b**, Experimental schema for co-labeling SNAP-H4 with TMR-Star (red) after BG-Nor1 (black) and DNA by EdU (green) in the same animal. **c**, Tissues from *Snap-H4c3* animals treated with CCl4 as indicated in (b) & imaged at 488 nm and 570 nm by confocal; scale bar 10 μm, imaged at 20x magnification. **d**, Mander’s correlation plot for TMR-star signal over EdU signal from (c), n=2 animals, * p<0.05 by Welch’s. **e**, Quantification of TMR & EdU signal from (c), n=2, one-way ANOVA, Brown Forsythe. **f & g**, Genome browser view of liver RNA-seq normalized signal across representative replication-dependent histone gene clusters on chromosome 13 after control versus CCl4 treatment.

Histone production depends on specific transcriptional and post-transcriptional processes to generate histone mRNAs that lack polyadenylation, which is unique for protein-coding genes. To determine if these canonical mRNA processing reactions were carried out after liver stress, we examined RNA-sequencing reads (**Fig. 3f,g**). We clearly identified short, highly abundant transcripts induced after CCl4 corresponding to properly processed histone mRNAs. This indicates that S-phase progression may not be strictly required to generate properly processed histone mRNA transcripts after liver stress.

Next, we sought to generalize these results beyond liver by examining stress responses in other organs. The chemotherapeutic drug doxorubicin is a known cardiotoxin. We tested whether cardiotoxic doses of doxorubicin could induce new histone production in the heart (**Fig. S5**). Old histones were blocked in *Snap-H4c3* mice, then doxorubicin was injected followed by pulse labeling with BG-Oregon Green (**Fig. S5a**). No new histone production was observed in heart. In contrast, strong histone production was again induced in the liver (**Fig. S5b**,**c**). We confirmed that both tissues had similar γ-H2AX induction after doxorubicin (**Fig. S5d**,**e**), consistent with its effects as a DNA-damaging agent. Thus, not all tissues respond to stress by inducing histone production.

Beyond testing other tissues, we also sought to confirm results with other histone genes. We generated a *Snap-H4c1* knock-in mouse (**Fig. S6a**), thereby appending a SNAP tag to a different histone H4 gene. Database searching indicated that *H4C1* expression was not as widespread compared to *H4C3*. Indeed, labeling of all histones with a single injection of TMR-Star showed SNAP-H4C1 expression was restricted to intestine and spleen, in contrast to the pervasive detection of SNAP-H4C3 in all tissues (**Fig. S6b**,**c**). Although *H4C1* was not expressed under baseline considtions in the liver, it was substantially induced after CCl4 stress (**Fig. S6d**,**e**,**f**). These data confirm that *in vivo* SNAP labeling is robust for different genes, allowing for the detection and quantitation of basal and induced protein expression.

### A signaling pathway to induce replication-independent histone production

Since the burst of histone synthesis in the liver did not appear to be strictly associated with S-phase progression, we sought to further characterize how histones were induced. Liver regeneration is initiated by a rapid priming phase, during which inflammatory cytokines, particularly TNFα and IL-6 released from Kupffer cells, prepare quiescent hepatocytes to respond to injury by activating immediate-early gene expression and regenerative signaling pathways (*23,24,25*). Given that TNFα is a principal initiator of this priming response, we investigated whether blocking TNFα signaling would attenuate histone induction (**Fig. 4a**). Administration of neutralizing antibodies to TNFα before CCl4 injection caused a strong reduction in SNAP-H4 signal after CCl4 treatment, compared to isotype control antibody (**Fig. 4b**). This implicates early inflammatory signaling as an important step in the induction of canonical histone synthesis following acute stress.

**Fig. 4:**
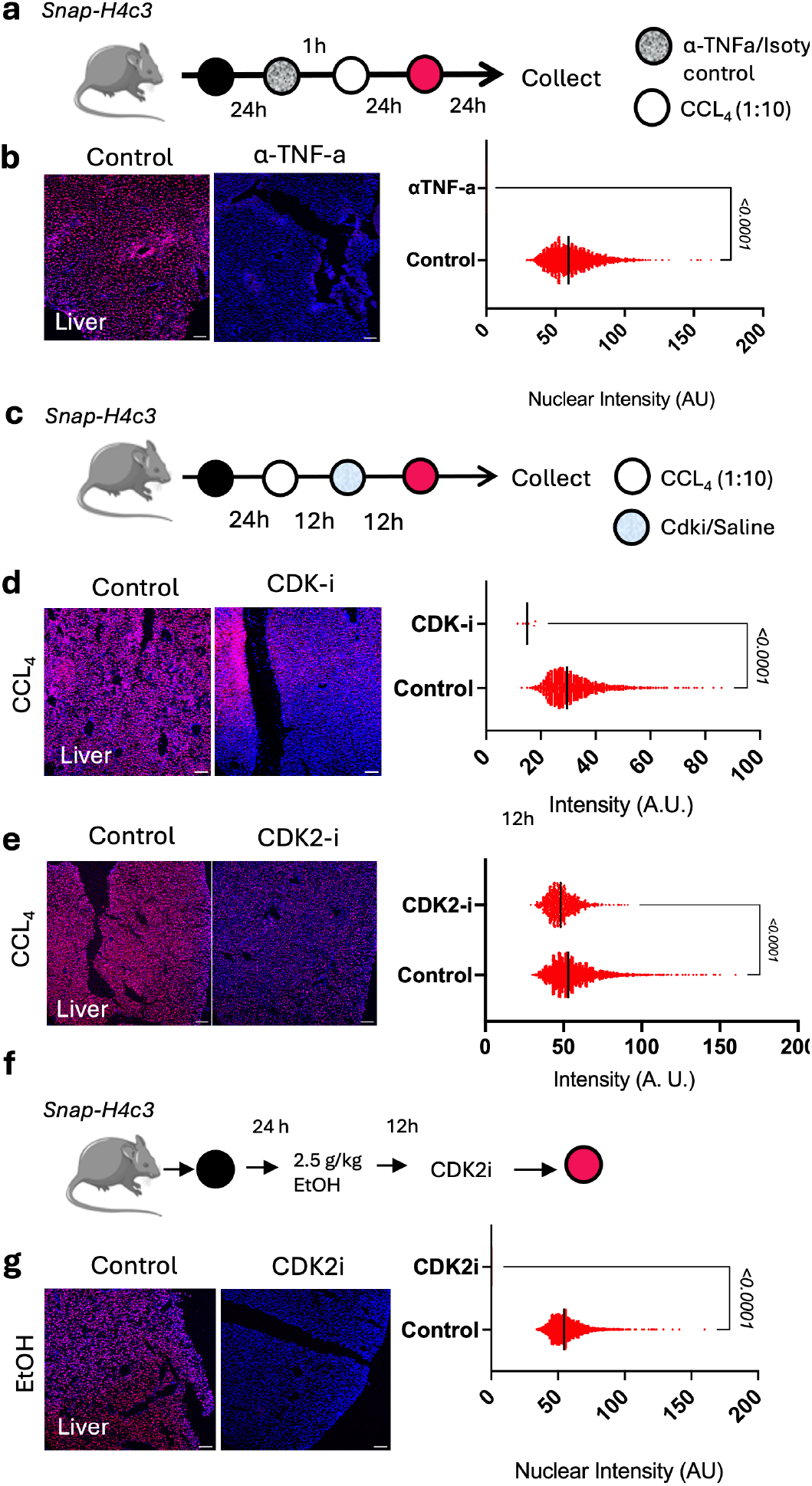
Blocking the histone surge. **a**, Schematic of experimental design where *Snap-H4c3* animals were treated with block, followed b4y0TNFαantibody and CCl4 (1:10), then injected with TMR-star before collection. **b**, Results of (a) where livers were harvested, flash frozen, cryo-sectioned, stained with DAPI & imaged at 570 nm by confocal, scale bar 10 μm, imaged at 10x magnification. Followed by quantification, n = 2, nuclear intensity per nuclei is plotted, column-based scatter plot (median), welch’s t-test. **c**, Schematic of *in vivo* CCl4 & CDK-inhibitor treatment in *Snap-H4*4*c*5*3* mice. **d**, Results of scheme (c); livers were harvested, flash frozen, cryo-sectioned, stained with DAPI & imaged at 570 nm by confocal, scale bar 10 μm, imaged at 10x magnification followed by quantification, n = 2, nuclear intensity per nuclei is plotted, column-based scatter plot (median), welch’s t-test. **e**, Results of *in vivo* CDK2i (roscovitine) treatment of *Snap-H4c3* animals; same procedure as (d). **f**, Schematic for ethanol (EtOH) treatment of *Snap-H4c3* animals, 2.5 g/kg binge for 12 hours followed by CDK2i trea5t0ment. **g**, Results of ethanol plus CDK2i treatment, same procedure as (d).

Kinase cascades downstream of TNFα converge on cyclin-dependent kinases (CDKs) to drive G1 entry. We tested whether blocking CDK activity could reverse histone production. First, we exposed *Snap-H4c3* MEFs to treatment with a CDK2 inhibitor (roscovitine) or a CDK4/6 inhibitor (palbociclib) or combination of each (**Fig. S7a**). Both inhibitors substantially reduced labeling of newly synthesized SNAP-H4, but the combination had the best reduction. Next, this was tested *in vivo* after liver injury. Mice were treated with BG-Nor1 to block old histones, then with CCl4, followed by combination of roscovitine plus palbociclib (CDK-i) versus vehicle, then TMR-Star labeling to detect newly synthesized histones (**Fig. 4c**). Compared to control, CDK-i treatment clearly reduced detection of SNAP-H4 signal (**Fig. 4d**). Treatment with CDK2 inhibitor (CDK2-i) alone had similar, but lesser effects (**Fig. 4e**). CDK-i effects were confirmed orthogonally by repeating the experiment in WT mice and pulse labeling newly synthesized histone with HPG methionine click analog. Again, CDK-i treatment strongly reduced labeling of newly synthesized histones (**Fig. S7b**). Also, ethanol-induced histone production was strongly abrogated by CDK2-i treatment (**Fig. 4f,g**). Thus, although stress-induced histone production occurs uncoupled from DNA synthesis, it still requires TNFα signaling, CDK2 and/or CDK4/6 activity.

### The chromatin impact of hepatocyte histone synthesis after stress

It is possible that widespread induction of core histone synthesis in hepatocytes after stress could contribute to epigenetic alterations required for response or adaptation. To characterize histone dynamics after hepatic stress, we treated *Snap-H4c3* mice with a single high dose of CCl4 and monitored new histone synthesis over time (**Fig. S7c**). Compared to the robust labeling of SNAP-H4 seen at 24 hours post-injury, the signal detected at 48 hours was markedly reduced (**Fig. S7d,e**), suggesting that histone production could be subsiding or excess histones were degraded. Next, to track the fate of pre-existing histones after injury, we labeled old SNAP-H4 with TMR-Star prior to CCl4 administration and harvested tissues at 24, 48, and 56 hours (**Fig. S7f**). We observed a progressive decline in TMR fluorescence over time (**Fig. S7g,h**), suggesting substantial loss or degradation of pre-existing histones following hepatic injury.

To more carefully define the behavior of pre-existing versus newly synthesized histones after CCl4-induced liver injury, we employed a dual-fluorophore pulse-chase labeling strategy (**Fig. S7i**). Pre-existing histones were first labeled with BG-Oregon Green, followed by CCl4 administration then labeling of newly synthesized histones with TMR-Star. In control animals, hepatocytes predominantly displayed green fluorescence, indicating retention of pre-existing histones (**Fig. S7j**). In contrast, CCl4-treated livers showed a marked loss of green signal accompanied by strong red fluorescence (**Fig. S7j**,**k**), suggesting replacement of old histones by newly synthesized ones.

To determine how inhibition of stress-induced histone dynamics affects the hepatic transcriptional response, we performed single-nuclei RNA sequencing (snRNA-seq) of liver tissue from control, CCl4-treated, CDKi-treated, and CCl4 + CDKi treated mice. Unsupervised clustering identified the major hepatic cell populations, including hepatocytes, Kupffer cells, neutrophils, monocytes, endothelial cells, cholangiocytes, fibroblasts, etc. (**Fig. 5a**). The distribution of nuclei across the UMAP was influenced by treatment condition, with CCl4 and CCl4 + CDKi samples exhibiting distinct transcriptional profiles relative to control animals (**Fig. 5b**). Further analyses focused solely on hepatocytes, which were the predominant cell type identified under all conditions (**Fig. 5c**).

**Fig. 5:**
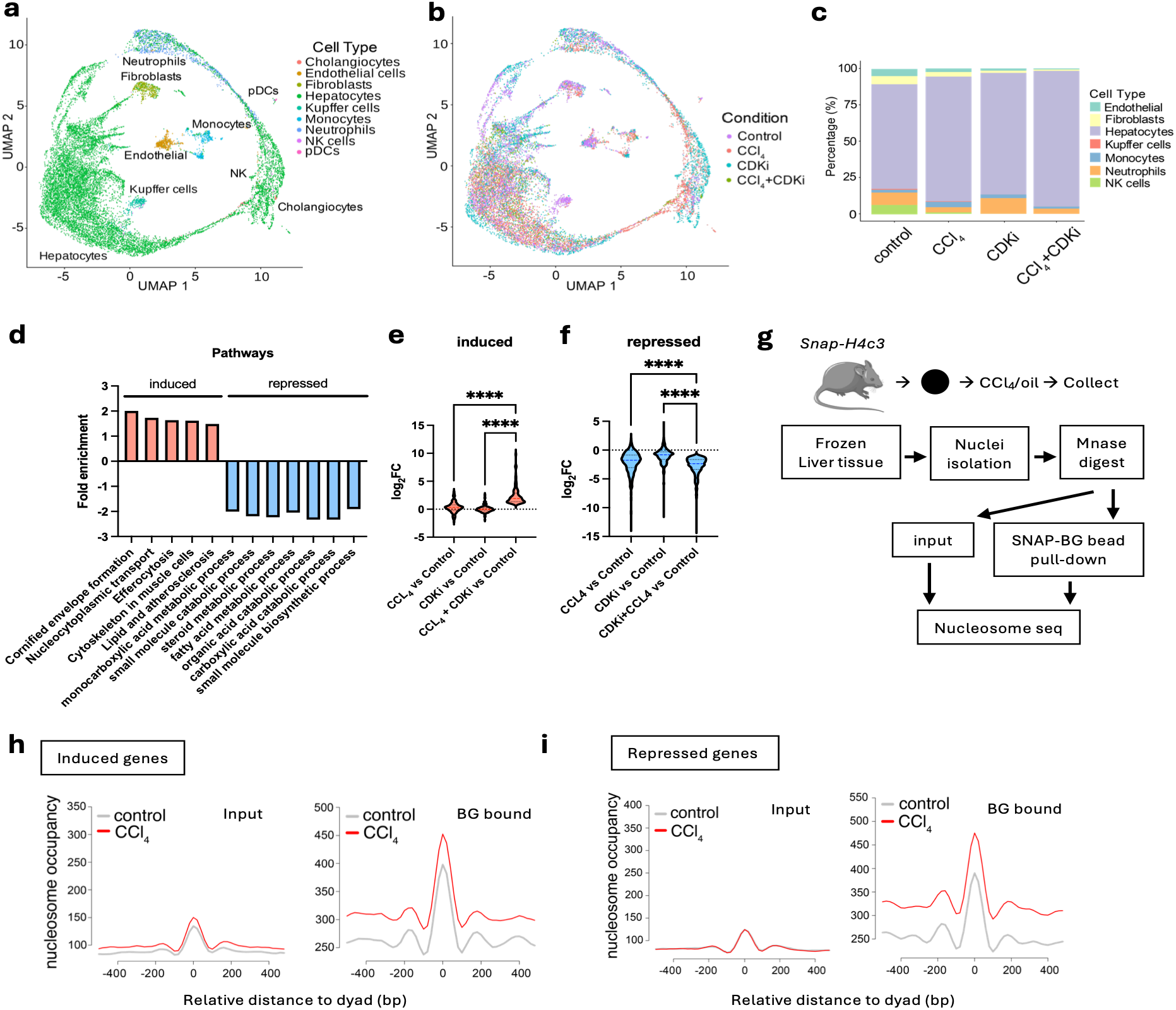
Gene expression buffering due to nucleosome incorporation after liver injury. **a**,**b**, UMAPs of snRNA-seq from livers treated with oil, CCl4, CDKi and combination (n = 3 each condition) showing clustering by cell type (a) and condition (b). **c**, Cell-type composition per condition. **d**, Pathway analysis of snRNA-seq genes induced or repressed by CCl4+CDKi versus control. **e**,**f**, violin plots showing log2FC gene expression across same gene sets for all conditions. n=187 induced and 447 reduced genes. **g**, Schematic of nucleosome-seq performed on CCl4 treated *Snap-H4c3* liver tissue. **h**,**i**, Aggregate dyad density profiles from nucleosome-seq libraries generated from *Snap-H4c3* mouse livers (n = 2 per condition) following CCl4 or control treatment. Metaplots show normalized dyad density centered on called nucleosome positions, stratified by snRNA-seq defined (h) induced genes, n=187, and (i) repressed genes, n=447. BG=benzylguanine. ****p<0.001 by ANOVA & Tukey’s.

Pathway analysis revealed that blocking new histone synthesis (CCl4+CDKi condition) induced pathways associated with cellular stress, while canonical hepatocyte metabolic programs were among the most strongly repressed programs (**Fig. 5d**). Importantly, inhibition of CDK activity after CCl4 treatment further altered the magnitude of these transcriptional responses compared to either CDKi or CCl4 treatment alone (**Fig. 5e,f**). Specifically, genes upregulated by CCl4 (n=187) were even more highly expressed after using CDKi to block new histone synthesis (**Fig. 5e**). Similarly, genes repressed after CCl4 treatment (n=447) were even further repressed after blocking new histone synthesis with CDKi (**Fig. 5f**). This suggested that new histone synthesis after liver stress could buffer gene expression, thereby limiting transcriptional changes induced by stress.

We sought to confirm that these gene expression changes were associated with new histone deposition at promoters of affected genes. Therefore, we used the *Snap-H4c3* mouse to trace incorporation of newly synthesized histones into chromatin of livers *in vivo. Snap-H4c3* mice were first treated with BG-Nor1 to irreversibly block all pre-existing (old) SNAP-H4 histones. Animals were next administered CCl4 or oil control for 24 hours to induce new histone synthesis, then liver nuclei were isolated. Extracted nuclei were subjected to MNase digestion followed by SNAP-affinity pull-down using BG-beads to enrich SNAP-H4-containing nucleosomes (**Fig. 5g**). Because all pre-existing SNAP-H4 had been blocked prior to injury, the recovered histones represent newly synthesized SNAP-H4 incorporated after treatment. Examination of nucleosome occupancy plots showed clear evidence of new histone deposition at both induced (n=187) and repressed (n=447) genes (**Fig. 5h, i**). Thus, new histones produced after liver stress are deposited at promoters of specific genes.

## Discussion

Histone synthesis has been difficult to assess in animals under physiological and stress conditions due to a lack of tractable measurement technologies. This work advances a molecular and chemical toolkit that allows for *in vivo* protein abundance monitoring in mammals. This allowed us to ascertain histone production across all tissues in a single pulse-chase experiment. As expected, under homeostatic conditions, histone production tracked closely with cell division rates in organs. However, following acute systemic and liver-specific stress, we observed an unexpected, coordinated induction of canonical core histone genes in the entire liver. These findings suggest acute organismal stress triggers a previously unknown histone synthesis program that operates broadly across hepatocytes and is not restricted to the subset of cells undergoing overt damage or DNA replication. This effect indicates that localized stress can stimulate an organ-wide response. We term this phenomenon ‘*histonekrexis,’* from the Greek *ékrēxis* (ἔκρηξις), meaning explosion, as the response is vigorous and quickly encompasses the entire liver.

The magnitude of *histonekrexis* raises the possibility that it could contribute to multiple aspects of liver homeostasis. Unlike most adult organs, the liver can rapidly restore mass and function following injury through coordinated transcriptional reprogramming and compensatory proliferation (*19*). The priming phase of liver regeneration is initiated by TNF-α and IL-6 secretion from Kupffer cells that stimulates a G0 to G1 transition in hepatocytes (*26*). Although bursts of histone synthesis in G1 may contribute to histone supply in preparation for S-phase (*27*), the data show that histone synthesis is uncoupled from DNA replication in majority of cells after stress.

We identify nucleosome deposition and transcriptional regulation as one consequence of new histone synthesis. As blocking *histonekrexis* accentuated transcriptional effects of liver stress, one important role of new histone production is to buffer gene expression, thereby stabilizing transcriptional output. This dovetails with prior work that nucleosome occupancy stabilizes gene expression fidelity (*28, 29, 30*), and influences regeneration (*31, 32*). In addition, recent work has identified a histone oxidoreductase activity (*33*), which may buffer against oxidation. Thus, in the setting of stress, a burst of histone synthesis via short, non-polyadenylated, non-spliced transcripts may be a parsimonious, stress-resilient adaptation to saturate chromatin with antioxidants. Similarly, in cancers S-phase independent histone synthesis may contribute to carcinogenesis (*34*) or therapeutic response (*35*).

In summary, this work demonstrates that histone synthesis is a dynamically regulated process in adult tissues. Instead of a mere consequence of DNA replication, various physiological and pathological stresses induce histone supply as an additional layer of chromatin regulation. Thus, dissecting the mechanisms that trigger *histonekrexis*, and determining how stress-induced histone synthesis influences gene expression and organismal response may inform recovery from tissue injury or inspire new therapies.

## Acknowledgments

The authors would like to acknowledge the following individuals for helpful feedback, discussions, training and administrative support during the course of this work: Skylar Nahi, Tripti Sharma, Xiuli Liu, Suman Komjeti, Prithvi Raj, Indu Raman, Chengsong Zhu, Noelle Williams. The following UTSW core facilities supported the work: Tissue Management Core Facility, CRI Mouse Genome Engineering Facility, Medicinal Chemistry Core, Microarray and Immune Phenotyping Core, Genomics Core, Preclinical Pharmacology Core.

## Author contributions

Conceptualization: SR, YW, MB, JJG

Methodology: SR, LV, JN, YW, JR

Investigation: SR, LV, JN, CG, HG, WKK, RG, SP

Funding acquisition: JJG

Project administration: MB, JJG

Supervision: SP, JR, MB, JJG

Writing – original draft: SR, JJG

## Data, code, and materials availability

Code and pipelines as well as source data used to generate figures for this work are available at the Gruber Lab github (https://github.com/GruberLabUTSW/SNAP-mouse).

## References and Notes

1. S.L. Commerford, A.L. Carsten, E.P. Cronkite. Histone turnover within nonproliferating cells. Proceedings of the National Academy of Sciences of the United States of America. 79, 1163–1165 (1982); doi:10.1073/pnas.79.4.1163.

2. J.A. Duerre, C.T. Lee. In vivo methylation and turnover of rat brain histones. Journal of neurochemistry. 23, 541–547 (1974). doi:10.1111/j.1471-4159.1974.tb06057

3. B.H. Toyama, J. N. Savas, S. K. Park, M. S. Harris, N. T. Ingolia, J. R. Yates, M. W. Hetzer. Identification of long-lived proteins reveals exceptional stability of essential cellular structures. Cell. 154, 971–982 (2013) doi:10.1016/j.cell.2013.07.037.

4. A. Groth, W. Rocha, A. Verreault, G. Almouzni. Chromatin Challenges during DNA Replication and Repair. Cell. 128, 721–733 (2007) doi:10.1016/j.cell.2007.01.030

5. M. D. Shmueli, D. Sheban, A. Eisenberg-Lerner, Y. Merbl, Histone degradation by the proteasome regulates chromatin and cellular plasticity. FEBS Journal. 289(12), 3304–3316 (2022). doi:10.1111/febs.15903

6. J. J. Gruber, B. Geller, A. M. Lipchik, J. Chen, A. A. Salahudeen, A. N. Ram, J. M. Ford, C. J. Kuo, M. P. Snyder. HAT1 Coordinates Histone Production and Acetylation via H4 Promoter Binding. Mol Cell. 75, 711–724 (2019) doi:10.1016/j.molcel.2019.05.034.

7. J. D. Gaddameedi, B. S. Geller, A. Rangarajan, T. A. Swaminathan, D. Dixon, K. Long, C. J. Golder, V. A. Vuong, S. Banuelos, R. Greenhouse, M. P. Snyder, A. M. Lipchik, J. J. Gruber, Acetyl-Click Screening Platform Identifies Small-Molecule Inhibitors of Histone Acetyltransferase 1 (HAT1). J. Med. Chem. 66, 5774–5801 (2023). doi:10.1021/acs.jmedchem.3c00039

8. J. Torné, D. Ray-Gallet, E. Boyarchuk, et al. Two HIRA-dependent pathways mediate H3.3 de novo deposition and recycling during transcription. Nat Struct Mol Biol 27, 1057–1068 (2020). doi:10.1038/s41594-020-0492-7

9. G. Yang, F. de Castro Reis, M. Sundukova, et al. Genetic targeting of chemical indicators in vivo. Nat Methods. 12, 137–139 (2015). doi:10.1038/nmeth.3207

10. B. D. Roger, et al. Genome-Wide Kinetics of Nucleosome Turnover Determined by Metabolic Labeling of Histones. Science 328, 1161–1164 (2010). doi:10.1126/science.1186777

11. Scharf AN, Barth TK, Imhof A. Establishment of histone modifications after chromatin assembly. Nucleic Acids Res. 2009. (15):5032–40. doi: 10.1093/nar/gkp518.

12. Zee BM, Levin RS, DiMaggio PA, Garcia BA. Global turnover of histone post-translational modifications and variants in human cells. Epigenetics Chromatin. 2010. 3(1):22. doi: 10.1186/1756-8935-3-22.

13. Alabert C, Barth TK, Reverón-Gómez N, Sidoli S, Schmidt A, Jensen ON, Imhof A, Groth A. Two distinct modes for propagation of histone PTMs across the cell cycle. Genes Dev. 2015 29(6):585–90. doi: 10.1101/gad.256354.114.

14. Arnaudo AM, Link AJ, Garcia BA. Bioorthogonal Chemistry for the Isolation and Study of Newly Synthesized Histones and Their Modifications. ACS Chem Biol. 2016. 11(3):782–91. doi: 10.1021/acschembio.5b00816.

15. A. Keppler, S. Gendreizig, T. Gronemeyer, et al. A general method for the covalent labeling of fusion proteins with small molecules in vivo. Nat Biotechnol 21, 86–89 (2003). doi:10.1038/nbt765

16. A. Gautier, A. Juillerat, C. Heinis, I. R. Corrêa, M. Kindermann, F. Beaufils, K. Johnsson, An Engineered Protein Tag for Multiprotein Labeling in Living Cells. Chemistry & Biology. 15, 128–136 (2008), doi:10.1016/j.chembiol.2008.01.007.

17. J. Wilhelm, S. Kühn, M. Tarnawski, et al. Kinetic and Structural Characterization of the Self-Labeling Protein Tags HaloTag7, SNAP-tag, and CLIP-tag. Biochemistry 60, 2560–2575 (2021). doi:10.1021/acs.biochem.1c00258

18. F. Coffey, B. Alabyev, T. Manser, Initial clonal expansion of germinal center B cells takes place at the perimeter of follicles. Immunity. 30, 599–609 (2009). doi:10.1016/j.immuni.2009.01.011.

19. A. M. Jamieson, P. Isnard, J. R. Dorfman, M. C. Coles, D. H. Raulet, Turnover and Proliferation of NK Cells in Steady State and Lymphopenic Conditions, The Journal of Immunology. 172, 864–870 (2004), doi:10.4049/jimmunol.172.2.864.

20. Y. Jia, L. Li, Y. Lin, et al. In vivo CRISPR screening identifies BAZ2 chromatin remodelers as druggable regulators of mammalian liver regeneration. Cell Stem Cell. 29, 372–385 (2022). doi:10.1016/j.stem.2022.01.001

21. Chembazhi UV, Bangru S, Dutta RK, Das D, Peiffer B, Natua S, Toohill K, Leona A, Purwar I, Bhowmik A, Goyal Y, Sun Z, Diehl AM, Kalsotra A. Dysregulated RNA splicing impairs regeneration in alcohol-associated liver disease. Nat Commun. 2025;16(1):8049. doi: 10.1038/s41467-025-63251-2.

22. Seemann S, Zohles F, Lupp A. Comprehensive comparison of three different animal models for systemic inflammation. J Biomed Sci. 2017;24(1):60. doi: 10.1186/s12929-017-0370-8.

23. Fausto N, Campbell JS, Riehle KJ. Liver regeneration. Hepatology. 2006;43(S1):S45–S53. doi:10.1002/hep.20969.

24. Tanimizu N, Miyajima A. Epithelial morphogenesis during liver development. Cold Spring Harb Perspect Biol. 2017;9(8):a027862. doi:10.1101/cshperspect.a027862.

25. Liu H, Zhang Y, Ning S. ScRNA-seq combined with ATAC-seq analysis to explore the metabolic balance mechanism of CCl4-induced liver inflammatory injury. Front Immunol. 2025; 16:1600685. doi: 10.3389/fimmu.2025.1600685.

26. Taub R. Liver regeneration: from myth to mechanism. Nat Rev Mol Cell Biol. 2004;5(10):836– 847. doi:10.1038/nrm1489.

27. Marzluff WF, Wagner EJ, Duronio RJ. Metabolism and regulation of canonical histone mRNAs: life without a poly(A) tail. Nat Rev Genet. 2008;9(11):843–843. doi:10.1038/nrg2438.

28. Wyrick JJ, Holstege FC, Jennings EG, Causton HC, Shore D, Grunstein M, Lander ES, Young RA. Chromosomal landscape of nucleosome-dependent gene expression and silencing in yeast. Nature. 1999. 402(6760):418–21. doi: 10.1038/46567.

29. Dreyer J, Ricci G, van den Berg J, Bhardwaj V, Funk J, Armstrong C, van Batenburg V, Sine C, VanInsberghe MA, Tjeerdsma RB, Marsman R, Mandemaker IK, di Sanzo S, Costantini J, Manzo SG, Biran A, Burny C, van Vugt MATM, Völker-Albert M, Groth A, Spencer SL, van Oudenaarden A, Mattiroli F. Acute multi-level response to defective de novo chromatin assembly in S-phase. Mol Cell. 2024. 84(24):4711-4728.e10. doi: 10.1016/j.molcel.2024.10.023. Epub 2024 Nov 12. Erratum in: Mol Cell. 2024 Dec 19;84(24):4945. doi: 10.1016/j.molcel.2024.11.036.

30. Kok JY, Harvey ZH, Axelsson E, Berger F. Nucleosome positioning shapes cryptic antisense transcription. PLoS Genet. 2026. 22(3):e1012078. doi: 10.1371/journal.pgen.1012078.

31. Cheloufi S, Elling U, Hopfgartner B, Jung YL, Murn J, Ninova M, Hubmann M, Badeaux AI, Euong Ang C, Tenen D, Wesche DJ, Abazova N, Hogue M, Tasdemir N, Brumbaugh J, Rathert P, Jude J, Ferrari F, Blanco A, Fellner M, Wenzel D, Zinner M, Vidal SE, Bell O, Stadtfeld M, Chang HY, Almouzni G, Lowe SW, Rinn J, Wernig M, Aravin A, Shi Y, Park PJ, Penninger JM, Zuber J, Hochedlinger K. The histone chaperone CAF-1 safeguards somatic cell identity. Nature. 2015. 528(7581):218–24. doi: 10.1038/nature15749.

32. Franklin R, Guo Y, He S, Chen M, Ji F, Zhou X, Frankhouser D, Do BT, Chiem C, Jang M, Blanco MA, Vander Heiden MG, Rockne RC, Ninova M, Sykes DB, Hochedlinger K, Lu R, Sadreyev RI, Murn J, Volk A, Cheloufi S. Regulation of chromatin accessibility by the histone chaperone CAF-1 sustains lineage fidelity. Nat Commun. 2022. 13(1):2350. doi: 10.1038/s41467-022-29730-6.

33. Attar N, Campos OA, Vogelauer M, Cheng C, Xue Y, Schmollinger S, Salwinski L, Mallipeddi NV, Boone BA, Yen L, Yang S, Zikovich S, Dardine J, Carey MF, Merchant SS, Kurdistani SK. The histone H3-H4 tetramer is a copper reductase enzyme. Science. 2020. 369(6499):59–64. doi: 10.1126/science.aba8740.

34. Henikoff S, Zheng Y, Paranal RM, Xu Y, Greene JE, Henikoff JG, Russell ZR, Szulzewsky F, Thirimanne HN, Kugel S, Holland EC, Ahmad K. RNA polymerase II at histone genes predicts outcome in human cancer. Science. 2025. 387(6735):737–743. doi: 10.1126/science.ads2169.

35. Moser SC, Khalizieva A, Roehsner J, Pottendorfer E, Kaptein ML, Ricci G, Bhardwaj V, Bleijerveld OB, Hoekman L, van der Heijden I, di Sanzo S, Fish A, Chikunova A, Haarhuis JHI, Oldenkamp R, Robbez-Masson L, Sprengers J, Vis DJ, Wessels LFA, van de Ven M, Pettitt SJ, Tutt ANJ, Lord CJ, Rowland BD, Völker-Albert M, Mattiroli F, Brummelkamp TR, Mazouzi A, Jonkers J. NASP modulates histone turnover to drive PARP inhibitor resistance. Nature. 2025. 645(8082):1071–1080. doi: 10.1038/s41586-025-09414-z.

36. Miura, H., R. M. Quadros, C. B. Gurumurthy, M. Ohtsuka, Easi-CRISPR for creating knock-in and conditional knockout mouse models using long ssDNA donors. Nat. Protoc. 13, 195–215 (2018). doi:10.1038/nprot.2017.153.

37. Feng, S., J. K. Laketa, J. H. Steinberg, A. T. Pypaert, M. H. Taunton, T. Balla, T. Inoue, A rapidly reversible chemical dimerizer system to study lipid signaling in living cells. Angew. Chem. Int. Ed. Engl. 53, 6720–6723 (2014). doi:10.1002/anie.201402766.

38. Sun, W.-C., K. R. Gee, D. H. Klaubert, R. P. Haugland, Synthesis of fluorinated fluoresceins. J. Org. Chem. 62, 6469–6475 (1997). doi:10.1021/jo970582k.

39. Srikun, D., A. E. Albers, C. I. Nam, A. T. Iavarone, C. J. Chang, Organelle-targetable fluorescent probes for imaging hydrogen peroxide in living cells via SNAP-tag protein labeling. J. Am. Chem. Soc. 132, 4455–4465 (2010). doi:10.1021/ja909931u

40. Ray-Gallet, D., A. Ricketts, K. Sato, R. Gupta, P. Boyarchuk, G. Almouzni, Two HIRA-dependent pathways mediate H3.3 de novo deposition and recycling during transcription. Nat. Struct. Mol. Biol. 18, 1249–1255 (2011). doi:10.1038/nsmb.2142.

41. N. Korenfeld, N. I. Toft, T. V. Dam, M. Charni-Natan, L. Grøntved, I. Goldstein, Protocol for bulk and single-nuclei chromatin accessibility quantification in mouse liver tissue. STAR Protoc. 4, 102462 (2023). doi:10.1016/j.xpro.2023.102462.

42. A. Forest, J.-P. Quivy, G. Almouzni, Mapping histone variant genomic distribution: Exploiting SNAP-tag labeling to follow the dynamics of incorporation of H3 variants. Methods Cell Biol. 182, 143–175 (2023). doi:10.1016/bs.mcb.2022.10.007.

