## Supplemental Methods and Figures for "Stress triggers global histone synthesis in the liver"

Materials and Methods

Genotyping strategy & generation of *Snap-H4c3* mouse

A SNAP-tag® coding sequence (Addgene #132427) was inserted in-frame at the N-terminus of the endogenous *Hist1h4c3* (H4C) locus to enable covalent labeling of histone H4 *in vivo*. The SNAP cassette was positioned immediately upstream of the *H4C3* coding sequence under the control of the native promoter, preserving endogenous regulation. *Snap-H4c3* transgenic line was generated using the *Easi*−CRISPR method as described previously (*36*). In brief, single guide RNAs (sgRNAs) were designed using the Broad Institute sgRNA design algorithm. Pure sgRNA (Synthego) was combined with recombinant Alt-R S.p. Cas9 nuclease V3 protein (Integrated DNA Technologies, 1081058) and was injected in C57BL/6J single-cell embryos with single-stranded DNA, containing SNAP tag flanked by homologous regions of the *H4C3* as the repair template. After microinjections, embryos were incubated overnight in a CO_2_ incubator at 37 °C. Surviving embryos were transplanted in pseudo-pregnant CD1 female mice to obtain live pups. Genotyping was performed using PCR on genomic DNA extracted from tail biopsies. Primer sets were designed to distinguish wild-type and knock-in alleles. Forward primer: GATGTCTGCTTTGTGGAATGG, Reverse primer (WT): GACAGTGGAAATCGTTTGTTAGTC Reverse primer (KI): CTTCACCACTTTCAGCAGTTTC. PCR amplification was performed using Platinum™ Direct PCR Universal Master Mix (Thermo Fisher Scientific) according to the manufacturer’s standard cycling parameters, only changing the annealing temperature (T_m_) according to primers specific to each allele. Amplicon sizes were 643bp for WT and 1186bp for knock-in alleles.

Heterozygous *Snap-H4c3* mice were used for all experiments.

Large-scale synthesis of BG-Oregon green, TMR-star and BG-Nor1

All reactions were performed with commercially obtained anhydrous solvents. All reactions and final products were monitored by liquid chromatography mass spectrometry (LC/MS) with an Agilent Technologies 1200 series LC/MS using electrospray ionization methods. Unless otherwise stated flash chromatography was performed with a Teledyne CombiFlash automated chromatography system. 1H and 13C NMR spectra were recorded on Varian Inova 400 MHz, Agilent 400 MHz, or Agilent 600 MHz spectrometer. Chemical shifts are reported relative to internal chloroform (CDCl3: 1H, = 7.26 ppm, 13C, = 77.36 ppm). Where spectra were recorded in mixtures of deuterated solvents, the more deuterated solvent was referenced. Coupling constants are in Hz and are reported as d (doublet), t (triplet), q (quartet), and m (multiplet).

Compounds **1**-**4** were synthesized by a modified procedure (*37*)

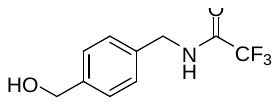

**2,2,2-trifluoro-N-(4-(hydroxymethyl)benzyl)acetamide (1).**

Triethylamine (6.4 ml, 45.9 mmol) was added in a rapid dropwise manner to a stirring solution of (4-(aminomethyl)phenyl)methanol (3.12 g, 22.8 mmol) in 104 ml of methanol. Ethyl trifluoroacetate (3.3 ml, 27.7 mmol) was then added dropwise to the reaction mixture, which was stirred overnight at ambient temperature. The reaction mixture was condensed, then taken up in EtOAc, washed twice with water and then brine, dried over Na_2_SO_4_, filtered and condensed to give 5.31 g solid (Yield=99%). The crude material was carried forward without further purification. ^1^H NMR (400 MHz, DMSO-*d*_6_) δ 9.98 (t, *J* = 6.1 Hz, 1H), 7.29 (d, *J* = 8.2 Hz, 2H), 7.22 (d, *J* = 8.1 Hz, 2H), 5.15 (t, *J* = 5.7 Hz, 1H), 4.47 (d, *J* = 5.7 Hz, 2H), 4.36 (d, *J* = 6.0 Hz, 2H). ESI-MS (*m/z*): 232.1 [M+H]^+^.

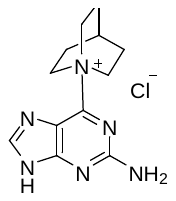

**6-(1l4-azabicyclo[2.2.2]octan-1-yl)-9H-purin-2-amine chloride (2).**

Triethylamine (1.05 ml, 7.6 mmol) was added to a solution of quinuclidine hydrogen chloride (1.11g, 7.6 mmol) in DMF (37 ml) and stirred for 30 minutes before 6-chloro-9H-purin-2-amine (1.3 g, 7.6 mmol) was added. The reaction was stirred overnight. Acetone was added to the heterogenous mixture before filtering. The white solid obtained was washed with acetone three times and air dried to give 1.33 g solid (Y=63%) which was carried forward without any further purification. ^1^H NMR (400 MHz, DMSO-*d*_6_) δ 8.34 (s, 1H), 7.09 (s, 2H), 4.23 – 4.10 (m, 6H), 2.24 (p, *J* = 3.3 Hz, 1H), 2.08 (dq, *J* = 8.3, 3.6 Hz, 6H). ESI-MS (*m/z*): 245.1 [M+H]^+^.

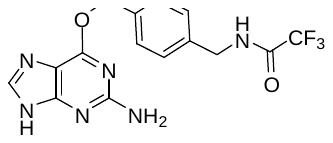

**N-(4-(((2-amino-9H-purin-6-yl)oxy)methyl)benzyl)-2,2,2-trifluoroacetamide (3).**

A solution of **1** (520 mg, 2.23 mmol) in 6.8 ml DMF was treated with sodium hydride (60% in mineral oil, 215 mg, 5.38 mmol) for 30 minutes before the addition of **2** (501 mg, 1.78 mmol) and a catalytic amount of DMAP. The reaction was heated overnight at 60 ^o^C. The cooled reaction was quenched with 0.5 ml H_2_O and condensed under high vacuum being heated to 40 ^O^C. The mixture was taken up in MeOH and adsorbed onto silica and dried under vacuum, before purification using automated chromatography over silica gel using 0-25% MeOH/DCM (+0.1% TEA). Isolated fractions collected at wavelength=220, 290 nm. Isolated mass=294.8 mg (Yield=45%). ^1^H NMR (600 MHz, DMSO-*d*_6_) δ 12.42 (s, 1H), 10.02 (t, *J* = 5.6 Hz, 1H), 7.81 (s, 1H), 7.49 (d, *J* = 7.9 Hz, 2H), 7.30 (d, *J* = 8.0 Hz, 2H), 6.30 (s, 2H), 5.46 (s, 2H), 4.39 (d, *J* = 4.4 Hz, 2H). ESI-MS (*m/z*): 367.2 [M+H]^+^.

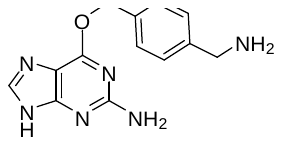

**6-((4-(aminomethyl)benzyl)oxy)-9H-purin-2-amine (4).**

A methylamine solution (33% in EtOH, 2 ml) was added to a solution of **3** (39.6 mg, 0.11 mmol) in MeOH (1ml). The reaction was stirred overnight, monitored by HPLC-MS, then condensed to an off-white solid and carried forward immediately. ESI-MS (*m/z*): 271.1 [M+H]^+^.

**BG-OREGON GREEN**

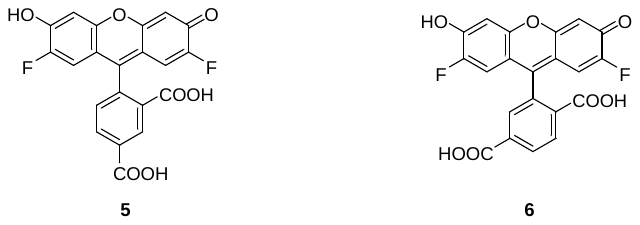

**Mixture of 4-(2,7-difluoro-6-hydroxy-3-oxo-3H-xanthen-9-yl)isophthalic acid (5)** and **isomer** **(6).**

Oregon green was synthesized using a modification of method (*38*). Trimellitic anhydride (750 mg, 3.90mmol) was added to a solution of 4-fluororesorcinol (1.0 g, 7.8 mmol) in 3.8 ml of methanesulfonic acid. The reaction was heated at 80 ^o^C overnight. The cooled reaction was poured over 30ml of ice water and the orange solid was filtered and washed with copious amounts of water. The solid was dried overnight in a vacuum oven at 60 ^o^C to give 1.97 g product which is a mixture of **5** and **6**. ^1^H NMR (400 MHz, DMSO-*d*_6_) δ 8.39 (dd, *J* = 1.5, 0.8 Hz, 1H), 8.29 (dd, *J* = 8.0, 1.5 Hz, 1H), 8.23 (dd, *J* = 8.0, 1.3 Hz, 1H), 8.10 (dd, *J* = 8.0, 0.8 Hz, 1H), 7.70 (t, *J* = 1.0 Hz, 1H), 7.41 (dd, *J* = 8.0, 0.8 Hz, 1H), 6.89 (d, *J* = 7.5 Hz, 4H), 6.62 (dd, *J* = 11.3, 9.6 Hz, 5H). ESI-MS (*m/z*): 413.2 [M+H]^+^.

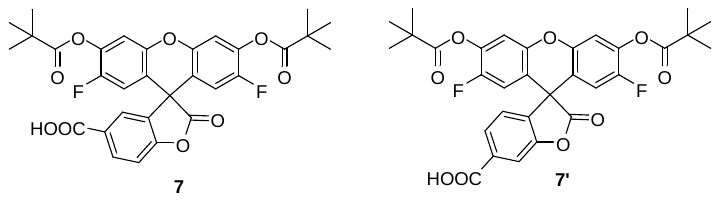

**2',7'-difluoro-2-oxo-3',6'-bis(pivaloyloxy)-2H-spiro[benzofuran-3,9'-xanthene]-6-carboxylic acid (7) and isomer 7’.**

Triethylamine (1.1 ml, 8.1 mmol) was added to a solution of **5** and **6** (1.5 g, 3.64 mmol) in DCM (35 ml). After 10 minutes of stirring, pivaloyl chloride (0.99 ml, 0.80 mmol) was added dropwise. The reaction was protected from light and stirred overnight. The reaction was condensed and purified twice via ISCO flash column chromatograph in 0-5% MeOH/DCM. The isolated desired product was washed with hot hexanes to remove traces of pivaloyl chloride/acid to give 360 mg of product as a mixture of isomers. Yield=17%. ^1^H NMR (400 MHz, Chloroform-*d*) δ 8.81 (dd, *J* = 1.5, 0.7 Hz, 1H), 8.44 (ddd, *J* = 15.3, 8.0, 1.4 Hz, 2H), 8.19 (dd, *J* = 7.9, 0.8 Hz, 1H), 7.92 (t, *J* = 1.0 Hz, 1H), 7.34 (dd, *J* = 8.1, 0.8 Hz, 1H), 7.16 (dd, *J* = 6.4, 3.7 Hz, 4H), 6.59 (dd, *J* = 9.7, 2.2 Hz, 4H), 1.40 (s, 40H). ESI-MS (*m/z*): 581.3 [M+H]^+^.

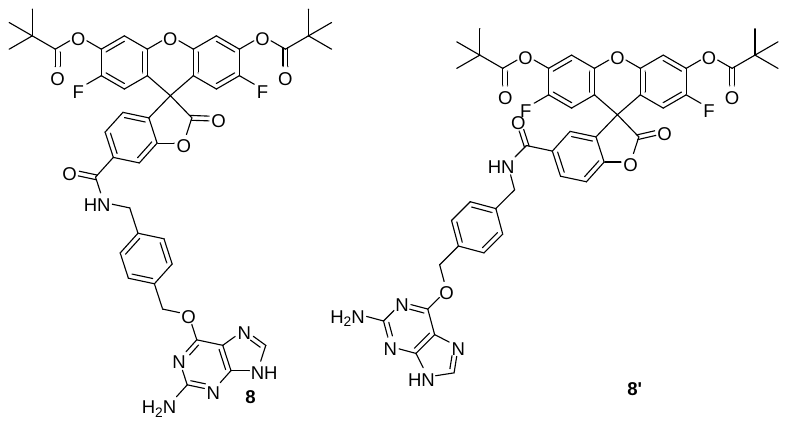

**6-((4-(((2-amino-9H-purin-6-yl)oxy)methyl)benzyl)carbamoyl)-2',7'-difluoro-2-oxo-2H-spiro[benzofuran-3,9'-xanthene]-3',6'-diyl bis(2,2-dimethylpropanoate) (8) and isomer 8’ (BG-Oregon Green).**

A solution of **7** and **7’** (133.3 mg, 0.23 mmol) in 1.6 ml DMF was pretreated with HATU (95.8 mg, 0.25 mmol) for 20 minutes followed by the addition of freshly prepared **4** (61.9 mg, 0.23 mmol) and DIPEA (40 uL, 0.23 mmol). The reaction had gone to completion within 4 hours. The mixture was diluted with EtOAc, and washed with 1M HCl, 4 x H_2_O, saturated NaHCO_3_ and then brine. The organic layer was dried over Na_2_SO_4_, filtered and condensed. The mixture was purified via ISCO flash column chromatography in 0-20% MeOH/DCM. Isolated 87 mg as a mixture of isomers (yield=45.5%). ^1^H NMR (600 MHz, DMSO-*d*_6_) δ 12.41 (d, *J* = 4.4 Hz, 2H), 9.45 (t, *J* = 5.9 Hz, 1H), 9.24 (q, *J* = 6.7, 6.0 Hz, 1H), 8.58 (dd, *J* = 1.7, 0.8 Hz, 1H), 8.34 (dd, *J* = 8.1, 1.6 Hz, 1H), 8.25 – 8.20 (m, 1H), 8.16 – 8.11 (m, 1H), 7.83 (t, *J* = 1.1 Hz, 1H), 7.79 (dd, *J* = 9.9, 0.6 Hz, 1H), 7.56 – 7.51 (m, 3H), 7.50 – 7.46 (m, 2H), 7.45 – 7.41 (m, 1H), 7.40 – 7.36 (m, 2H), 7.31 – 7.28 (m, 1H), 7.03 (dd, *J* = 10.3, 1.6 Hz, 3H), 6.29 (d, *J* = 10.9 Hz, 3H), 5.44 (d, *J* = 25.4 Hz, 3H), 4.54 (d, *J* = 5.8 Hz, 2H), 4.43 (d, *J* = 5.8 Hz, 2H), 4.11 (q, *J* = 5.3 Hz, 1H), 3.17 (d, *J* = 5.3 Hz, 2H), 1.32 (d, *J* = 2.0 Hz, 26H). ESI-MS (*m/z*): 833.2, [M+H]^+^.

**BG-NOR1**

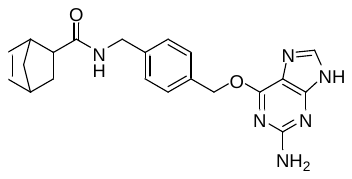

**N-(4-(((2-amino-9H-purin-6-yl)oxy)methyl)benzyl)bicyclo[2.2.1]hept-5-ene-2-carboxamide(9)**

Solid 3-(((ethylimino)methylene)amino)-N,N-dimethylpropan-1-amine hydrochloride (EDC, 64.4mg, 0.34 mmol), 1H-benzo[d][1,2,3]triazol-1-ol hydrate (HOBT, 42.2 mg, 0.28 mmol) and bicyclo[2.2.1]hept-5-ene-2-carboxylic acid (89.0 mg, 0.64 mmol) were added to a vial containing **4** (73.0 mg, 0.24 mmol). DMF (2.6 ml) was added, followed by trimethylamine (49 ul, 0.35 mmol). After overnight stirring conversion was low with mostly **4** remained. A microspatula full of waxy bicyclo[2.2.1]hept-5-ene-2-carboxylic acid was added and stirred for 6 days. The reaction was stopped despite low conversion. The reaction was diluted with EtOAc and washed several times with H_2_O and then brine. The organic layer was dried over Na_2_SO_4_, filtered and condensed. The mixture was purified via ISCO FCC in 0-10% MeOH/DCM(+0.1% Et_3_N). Isolated 12.5 mg in 12% yield. ^1^H NMR (400 MHz, Methanol-*d*_4_) δ 7.70 (s, 1H), 7.41 (d, *J* = 7.8 Hz, 2H), 7.21 (d, *J* = 7.8 Hz, 2H), 6.15 (dd, *J* = 5.8, 3.1 Hz, 1H), 5.87 (dd, *J* = 5.7, 2.8 Hz, 1H), 5.48 (s, 2H), 4.41 – 4.24 (m, 2H), 3.16 – 3.11 (m, 1H), 2.91 (dt, *J* = 9.3, 4.0 Hz, 1H), 2.88 – 2.83 (m, 1H), 1.87 (ddd, *J* = 11.5, 9.3, 3.7 Hz, 1H), 1.43 – 1.32 (m, 2H), 1.31 – 1.21 (m, 1H). ESI-MS (*m/z*): 391.2 [M+H]^+^.

**TMR-STAR**

Intermediates **10** and **11** were synthesized following a procedure from (*39*).

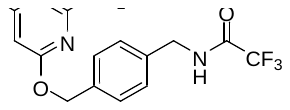

**N-(4-(((2-amino-6-chloropyrimidin-4-yl)oxy)methyl)benzyl)-2,2,2-trifluoroacetamide (10).**

A solution of 1 (301.5 mg, 1.29 mmol) in 4.3 ml DMAC was cooled to -20 ^o^C in a chiller before the addition of sodium hydride (60% dispersion in mineral oil, 135 mg, 3.38 mmol). The mixture was stirred cold for 30 minutes before the solution of 4,6-dichloropyrimidin-2-amine (296.6 mg, 1.81 mmol) in 6.4 ml DMF was added dropwise. The mixture was gently warmed to ambient temperature and stirred overnight. The reaction mixture was diluted with EtOAc, washed several times with H_2_O and then brine. The organic layer was dried over Na_2_SO_4_, filtered and condensed. The crude mixture was purified by automated flash chromatography in 0-100% EtOAc/DCM to give 315 mg of desired compound in 76.5% yield. ^1^H NMR (600 MHz, DMSO-*d*_6_) δ 10.02 (s, 1H), 7.44 – 7.40 (m, 2H), 7.30 – 7.26 (m, 2H), 7.12 (s, 2H), 6.14 (s, 1H), 5.29 (s, 2H), 4.38 (s, 2H). ESI-MS (*m/z*): 361.1 [M+H]^+^.

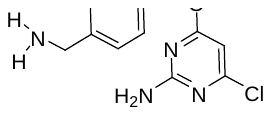

**4-((4-(aminomethyl)benzyl)oxy)-6-chloropyrimidin-2-amine(11).**

Methanamine (4 ml) was added to a solution of **10** (71.0 mg, 0.20 mmol) in 2 ml MeOH. The reaction was stirred overnight, then monitored by HPLC-MS until complete. The reaction was condensed under reduced pressure and carried forward immediately. ESI-MS (*m/z*): 265.1 [M+H]^+^.

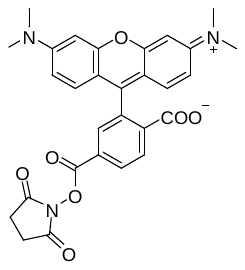

**2-(6-(dimethylamino)-3-(dimethyliminio)-9,9a-dihydro-3H-xanthen-9-yl)-4-(((2,5-dioxopyrrolidin-1-yl)oxy)carbonyl)benzoate(12)**

N,N-dimethylpyridin-4-amine (DMAP, 56.4 mg, 0.46 mmol) was added to an oven dried flask containing 5-carboxytetramethylrhodamine (100.4 mg, 0.23mmol). DMF, 2 ml and Hunig’s base (40.5 uL, 0.23 mmol) were added followed by slow portionwise addition of disuccinimyl carbonate (96.0 mg, 0.38 mmol). After 4 hours of stirring protected from light, about 85% conversion to the succinimide was achieved. Charging more disuccinimyl carbonate and/or longer reaction times did not increase conversion. Cold Et_2_O (20 ml) was added to the reaction mixture, which was then sonicated and maintained on ice to settle the purple precipitate. The filtrate was pipetted out and the mixture washed another 2x with cold Et_2_O, significantly removing most of the DMAP and some unreacted 5-carboxytetramethylrhodamine and little desired. The purple solid was dried under vacuum before being taken directly into the acylation. ESI-MS (*m/z*): 528.2 [M+H]^+^.

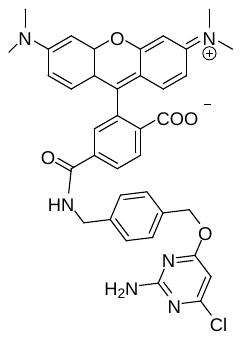

**4-((4-(((2-amino-6-chloropyrimidin-4-yl)oxy)methyl)benzyl)carbamoyl)-2-(6-(dimethylamino)-3-(dimethyliminio)-8a,9,9a,10a-tetrahydro-3H-xanthen-9-yl)benzoate (12) (SNAP-Cell TMR-Star)**

A solution of amine **11** (53.8 mg, 0.42 mmol) in 2 ml DMF was added to succinimde **12** (Assumed 110 mg, 0.21 mmol) followed by addition of iPr_2_NEt (72.5 ul, 0.42 mmol). The reaction was stirred overnight protected from light. The reaction mixture was diluted with EtOAc, and washed 3 x saturated NaHCO_3_.The organic layer was dried, then loaded in MeOH and condensed. The mixture was purified via ISCO reverse phase chromatography in 0-40% ACN/H_2_O(+0.1% acetic acid). Isolated 26.3 mg in 19% yield. ^1^H NMR (600 MHz, DMSO-*d*_6_) δ 9.25 (t, *J* = 6.0 Hz, 1H), 8.20 (dd, *J* = 8.1, 1.4 Hz, 1H), 8.07 (dd, *J* = 8.1, 0.7 Hz, 1H), 7.68 – 7.65 (m, 1H), 7.37 – 7.32 (m, 2H), 7.28 – 7.24 (m, 2H), 7.10 (s, 2H), 6.55 – 6.46 (m, 6H), 6.10 (s, 1H), 5.25 (s, 2H), 4.40 (d, *J* = 5.9 Hz, 2H), 2.94 (s, 12H). . ESI-MS (*m/z*): 677.2 [M+H]^+^.

Pharmacokinetics of TMR-star and BG-Nor1

Pharmacokinetic studies were performed to establish dosing regimens for *in vivo* SNAP-tag blocking and labeling experiments. Male and female mice (wild-type C57BL/6J) received a single intraperitoneal (i.p.) administration of either BG-Nor1 or TMR-Star at 10 mg/kg. Both compounds were formulated at 1 mg/mL in a vehicle consisting of 10% DMSO, 10% Cremophor EL (Sigma, C5135), and 80% Dextrose [5%] water (D5W). Injection volumes ranged from 0.16–0.22 mL depending on animal body weight. Plasma samples were collected from three mice per timepoint at 0, 10, 30, 90, 180, 360, 960, and 1440 minutes by submandibular bleeding.

Plasma concentrations of BG-Nor1 and TMR-Star were quantified by LC-MS/MS using a Sciex 4500 triple quadrupole mass spectrometer operated in positive electrospray ionization (ESI) mode with multiple reaction monitoring (MRM). Chromatographic separation was performed using an Agilent Poroshell 120 EC-C18 column (50 × 3.0 mm, 2.7 μm particle size). Mobile phase A consisted of water containing 0.1% formic acid, and mobile phase B consisted of methanol containing 0.1% formic acid. For BG-Nor1 analysis, the gradient was 0–0.5 min, 97% A; 0.5–1.0 min, gradient to 100% B; 1.0–3.5 min, 100% B; 3.5–3.6 min, gradient to 97% A; 3.6–4.5 min, 97% A. For TMR-Star analysis, the gradient was 0–1.0 min, 97% A; 1.0–2.0 min, gradient to 100% B; 2.0–3.5 min, 100% B; 3.5–3.6 min, gradient to 97% A; 3.6–4.5 min, 97% A. The flow rate was maintained at 0.8 mL/min. Injection volumes were 5 μL for BG-Nor1 and 10 μL for TMR-Star analyses.

For sample preparation, 98 μL of blank mouse plasma was combined with 2 μL of standard or quality-control stock solution and mixed with 200 μL of ice-cold acetonitrile containing 0.15% formic acid and an internal standard (final concentration 50 ng/mL). Samples were vortexed for 15 seconds, incubated at room temperature for 10 minutes, and centrifuged at 13,200 rpm for 3 minutes. Supernatants were transferred to LC-MS/MS vials for analysis. BG-Nor1 and TMR-Star were detected using compound-specific MRM transitions of m/z 391.296→174.000 and m/z 677.153→518.100, respectively. Pharmacokinetic parameters were calculated by noncompartmental analysis using Phoenix WinNonlin (Certara)(Fig.S2b, 2c). These pharmacokinetic properties informed the design of subsequent *in vivo* pulse-chase labeling experiments.

Isolation and immortalization of MEFs

Mouse embryonic fibroblasts (MEFs) were isolated from E12 embryos obtained from timed pregnancies. Embryos were dissected; head and internal organs were removed. Remaining tissue was minced and digested in collagenase solution at 37°C. Cells were plated in DMEM (Sigma D6429) supplemented with 10% fetal bovine serum (FBS) (biowest, S1620, 297H23) and 1% penicillin/streptomycin(P/S) (Gibco, 15-140-122). Primary MEFs were expanded for 2 passages, the fast-growing cells were snap frozen for further use. Some were left to expand by repeatedly culturing them every 3 days (spontaneous immortalization) for >10 weeks. Successfully immortalized MEFs were maintained under standard culture conditions.

*In vitro* SNAP-MEFs assay

Immortalized *Snap-H4c3* MEFs were maintained in DMEM supplemented with 10% FBS and 1% P/S at 37°C with 5% CO₂. Cells were plated on coverslips in 24-well plates and allowed to adhere for 24-36 h prior to SNAP labeling. For SNAP labeling, culture medium was removed and replaced with fresh complete medium containing TMR-star (2mM stock in DMSO) at a final concentration of 2 µM in 200 µL per well. Cells were incubated at 37°C for 30 min. Following labeling, cells were washed once with 1x PBS and incubated in fresh dye-free media for an additional 30 min at 37°C to allow diffusion and removal of excess unbound substrate. For fixation, cells were incubated in 4% paraformaldehyde (PFA) (250 µL per well) for 15–20 min at room temperature (RT). PFA was removed completely, and coverslips were washed twice with 1x PBS. Then 250 uL of 0.2% Triton-X was added to wells, incubated at RT for 5 mins, then washed thrice with Mol Bio (molecular biology) grade water of which final wash was with 1x DAPI. Coverslips were mounted using Immu-Mount (epredia) and sealed prior to imaging. For blocking experiments, cells were pre-incubated with SNAP-Cell^®^ Block or BG-Nor1 (in-house) substrate 24 h before TMR-Star labeling using the same labeling and washout conditions. This assay was modified from previously described SNAP-tag labeling method (*40*).

Nuclei isolation from liver tissue

For all nuclei isolation, frozen tissues were used. The STAR protocol (*41*) was used with some modifications: we did not use spermidine in the hypotonic buffer, and the nuclei pellet was counted with trypan blue & snap frozen for further assays.

*In vivo* SNAP labeling

For *in vivo* labeling of SNAP-H4, mice were administered SNAP substrates via i.p. injection. TMR-Star was used to label existing SNAP-H4, while BG-Nor1 was used for blocking pre-existing SNAP sites. For pulse-chase experiments, mice were injected with BG-Nor1 (10 mg/kg) to block existing SNAP sites, followed by TMR-Star (5 mg/kg) after 24 or 48 h. Mice were sacrificed 24 h after TMR-Star injection. For single-label experiments, TMR-Star (5 mg/kg) was administered, and tissues were harvested at indicated time points. Organs were collected, flash frozen, & stored in -80C until further use.

Cryosectioning and imaging

The frozen tissues were embedded in OCT (Leica Surgipath FSC 22), and cryosectioned at thickness of 10 µm using Leica cryostat (CM3050 S). Sections were left to dry overnight at RT, then directly mounted with glass coverslips using Vectashield antifade mounting medium with DAPI (Vector labs), then sealed. Imaging was performed using a ZEISS 700 confocal microscope equipped with appropriate filters for TMR-star (excitation ~554 nm, emission ~580 nm). Exposure settings were kept constant across conditions.

Image processing using FIJI (ImageJ)

Raw Zeiss microscopy files (.czi) were processed using FIJI/ImageJ (v2.16.0 / 1.54g / Java 1.8.0_322 (64-bit)). Images were imported into FIJI using the Bio-Formats importer with channel splitting enabled. Individual fluorescence channels were separated, and display thresholds were adjusted using Image → Adjust → Threshold. For each experimental group, identical threshold values were applied to all images within a given channel to ensure consistent visualization and comparison across samples.

Following threshold adjustment, channels were recombined using Image → Color → Merge Channels, with DAPI assigned to the blue channel and TMR-Star assigned to the red channel. Additional channels were assigned as appropriate for each experiment. Scale bars were added using Analyze → Tools → Scale Bar, with scale parameters determined from the microscope metadata associated with each image.

Processed images were exported as TIFF files for archival and quantitative analysis. Then as PNG files for figure preparation and manuscript presentation. No image-specific adjustments were performed beyond uniform thresholding and channel assignment within individual experiments.

Signal quantification using CellProfiler

Fluorescence images were analyzed using CellProfiler (v4.2.8) using a custom pipeline developed for quantification of nuclear SNAP-H4 labeling. Multi-channel TIFF images were imported into CellProfiler and filtered using the ‘Images’ module to include image files only. Metadata were extracted from image filenames using the ‘Metadata’ module and used to assign image channels and experimental groups through the ‘NamesAndTypes’ and ‘Groups’ modules.

Nuclei were identified from the DAPI channel using the ‘IdentifyPrimaryObjects’ module. Object identification was performed using a Global Otsu thresholding strategy with two intensity classes. The expected object diameter range was set to 10-25 pixels to accurately segment individual nuclei while minimizing inclusion of debris and merged objects. Identified nuclei were used as the primary objects for all downstream measurements. To restrict quantification to nuclear regions, the identified nuclear masks were applied to the TMR-Star fluorescence channel using the ‘MaskImage’ module. Images were subsequently processed using the ‘EnhanceEdges’ module with Canny edge detection to improve object boundary definition and facilitate robust fluorescence measurements. Fluorescence intensity was quantified using the ‘MeasureObjectIntensity’ module.

Integrated fluorescence intensity within each segmented nucleus was used as the primary metric for quantification. In selected analyses, the total number of fluorescent objects identified within a field or tissue section was also recorded as an additional measure of labeling frequency. Single-cell measurements and image-level summary statistics were exported using the ‘ExportToSpreadsheet’ module and analyzed in GraphPad Prism.

Immunohistochemistry of Ki67and gH2AX

Liver and heart tissues were fixed in 4% PFA, paraffin embedded, and sectioned at a thickness of 4 μm. Tissue sections were deparaffinized in xylene and rehydrated through a graded ethanol series. Antigen retrieval was performed by heating sections in citrate buffer, pH 6.0 / EDTA buffer, pH 9.0. Endogenous peroxidase activity was quenched with 3% hydrogen peroxide, followed by blocking. Sections were incubated overnight at 4°C with primary antibodies against Ki67 (CST 12202, 1:400) or γH2AX (CST 9718, 1:200). Following washing, sections were incubated with an appropriate HRP-conjugated secondary antibody and developed using 3,3′-diaminobenzidine (DAB). Sections were counterstained with hematoxylin, dehydrated, cleared, and mounted. Images were acquired using Leica DM6000 B under identical imaging conditions. The percentage of Ki67- or γH2AX-positive staining was quantified from representative fields using ImageJ/Fiji and expressed as the percentage of positively stained area relative to the total tissue area.

Acid extraction of MEFs and liver tissue

Total histones were isolated by acid extraction. Cells from culture were resuspended in lysis buffer (0.5N HCl, 10% glycerol) and incubated on ice for 30 mins to extract histones. Samples were centrifuged to remove debris, and supernatants containing histones were collected. 100% cold acetone (1:1) was added to supernatant along with 20uL of mix [0.5M NaOH + 0.5M EDTA]. Histones were left to precipitate overnight at -20 or -80C. Extracted histones were quantified and used for SDS–PAGE and downstream analyses.

For liver tissue, first nuclei were isolated using previously described methods. Then the nuclei pellet was subject to the same above acid extraction.

Tissue collection and western blots

Mice were euthanized at indicated time points, and tissues were rapidly harvested and snap-frozen in liquid nitrogen. For liver injury experiments, mice were treated with CCl₄ diluted at 1:10 (~1.5 mg/kg) in peanut oil and analyzed after 24 or 48 hours or as specified. Tissues were homogenized using a cold mortar & pestle, in lysis buffer (1x RIPA) supplemented with protease inhibitors. The crude mix was transferred to 1.5mL tube in ice. The sample were subjected to ultrasonication (10sec, 10% power). Then clarified by spinning at 15,000x g for 5 min, 4°C. Protein concentration was determined using BCA assay. Equal amounts of protein were resolved on SDS–PAGE gels and transferred to nitrocellulose membranes. Membranes were blocked in 5% milk in PBST and incubated with primary antibodies overnight at 4°C. Primary antibodies included anti-histone H3, H4 (CST #9715, #2592), histone modification antibodies H3K27me3, H3K9ac (CST #9733, ab218553) and anti-SNAP (NEB, #P9310S). Following incubation with HRP-conjugated secondary antibodies (CST #7074S), signals were detected using chemiluminescence.

*In vivo* EdU labeling

To assess DNA synthesis *in vivo*, WT mice were administered Oil or CCl₄ and allowed to recover for 24 h, after which 5-ethynyl-2′-deoxyuridine (EdU, Sigma #900584) was administered at dose of 5 mg/kg. After 4 h, the animals were sacrificed, liver & intestinal tissue were harvested and snap frozen. The tissues were sectioned, fixed on slides, then processed for EdU detection using a copper-catalyzed azide-alkyne cycloaddition reaction with fluorescent azide probes according to the manufacturer’s instructions (EdU Click-iT™, Thermo Fisher Scientific, C10337). Then the slides were covered with Vectashield mount, sealed, imaged and quantified on ZEISS 700 confocal microscope using Alexa488 filters.

For the detection of DNA synthesis plus SNAP-H4 in the same tissues, the *Snap-H4c3* mice were administered BG-nor1, after 24 h they were subject to oil or 1:10 CCl4, then after 6 h, 5 mg/kg EdU at repeated intervals (every 6 h) for a total of 24 h. Then SNAP-H4 labeling with TMR-Star was performed in parallel (as indicated in the experimental schematic, Fig.3c) to enable comparison of histone synthesis and DNA replication. At the indicated time point, mice were euthanized and tissues were harvested, & flash frozen. For sectioning & imaging the same process outlined above. For co-localization studies, TMR-Star signal was imaged directly without additional staining. Fluorescence imaging was performed using a ZEISS 700 confocal microscope with identical acquisition settings across conditions. Quantification of EdU and TMR-Star co-localization was performed using Manders’ overlap coefficient calculated in ImageJ, with thresholds applied uniformly across samples.

HPG-click assay with liver tissues

To assess nascent protein synthesis *in vivo*, mice were administered the methionine analog L-homopropargylglycine (Click-iT™ HPG, Thermo Fisher Scientific, C10186), via i.p. injection at a dose of 10 mg/kg and allowed to incorporate for 18 h (a total of 24 h after CCl_4_ treatment). Liver tissues were harvested, and snap frozen. Tissue samples were homogenized in ice-cold 1x RIPA buffer supplemented with protease inhibitors. Lysates were clarified by centrifugation at 10,000 × g for 10 min at 4°C, and protein concentration was determined using a BCA assay. For detection of nascent proteins, 20 µg of protein were subjected to copper(I)-catalyzed azide-alkyne cycloaddition using a biotin-azide probe (Click-iT™ Protein Reaction Buffer Kit, Thermo Fisher Scientific, C10276) according to the manufacturer’s instructions. Following the click reaction, samples were mixed with SDS loading buffer and resolved by SDS–PAGE. Proteins were transferred to nitrocellulose membranes. Biotin-labeled nascent proteins were detected using HRP-conjugated streptavidin (1:10,000 dilution), followed by chemiluminescent detection. To assess total protein loading, membranes were stained with memCode blue.

Sample prep for Nucleosome seq

*Snap-H4c3* mice were administered 10 mg/kg BG-Nor1 by i.p. injection to covalently block SNAP-H4. After 24 h, mice were treated with either CCl₄ or peanut oil, and livers were collected 24 h later. Liver tissue was rapidly harvested and snap frozen. Nuclei were isolated from liver tissue according to previously described methods (*41*). Chromatin was digested with micrococcal nuclease (MNase) under optimized conditions. Digestion was stopped with EDTA, and soluble nucleosomes were recovered after centrifugation. A fraction of this material was retained as input. SNAP-H4-containing nucleosomes were captured using benzylguanine-conjugated (BG) magnetic beads, adapted from the SNAP-capture protocol (*47*). Soluble mononucleosomes were incubated with BG-magnetic beads overnight at 4°C with gentle rotation to allow covalent capture of SNAP-tagged nucleosomes. Beads were washed extensively to remove nonspecifically bound chromatin. Captured nucleosomes were eluted by proteinase K/SDS digestion, and associated DNA was purified from both input and SNAP-captured fractions. DNA fragment size was assessed by TapeStation to confirm enrichment of mononucleosomal DNA. Sequencing libraries were prepared from purified DNA using NEB Multiplex Oligos kit (#E6440) and NEBNext Ultra II DNA kit (#E7645S) and sequenced on Illumina platform NovaSeq X using 150 bp paired-end reads (PE150).

Sample prep for bulk RNA-seq, rRNA depletion RNA-seq and single-nuclei RNA-seq

To assess whether introduction of the *Snap-H4c3* allele altered basal liver gene expression, bulk RNA sequencing was performed on liver nuclei isolated from untreated WT and *Snap-H4c3* mice (n = 3 each). Following isolation, 5 × 10^5 nuclei were transferred into DNA LoBind^®^ Tubes (Eppendorf, #022431021) and resuspended in 100 μL DNA/RNA Shield (Zymo Research) for nucleic acid preservation. Samples were submitted to Plasmidsaurus (San Francisco, CA) for RNA sequencing. RNA extraction, library preparation, and sequencing were performed by Plasmidsaurus according to their standard RNA-seq workflow.

For rRNA depletion RNA-seq, total RNA was extracted from liver tissue using TRIZOL, and further purified by chloroform, after which samples were run through Qiagen RNeasy Mini Spin Columns. Then the eluted sample was treated with DNase-I. RNA integrity was assessed using Qubit hs-RNA and bioanalyzer. Directional library preparation (rRNA removal) was performed by Novogene USA. Sequencing was performed on an Illumina platform NovaSeq X Plus series (PE150).

For single-nuclei RNA-seq, the nuclei from frozen liver tissues were isolated using methods defined earlier. The integrity of the nuclei was assessed using the automated LUNA-FL™ Dual Fluorescence Cell Counter. 100,000 nuclei that yields a coverage of 600-1000 nuclei was taken for cDNA prep using Chromium GEM-X Single Cell 3' Reagent Kits v4. Then the 10x Genomics single-nuclei 3’ gene expression libraries were prepared. Finally, they were sequenced on illumina NovaSeq X Sequencer.

Bioinformatics analysis methods

Bulk RNA-seq

For Plasmidsaurus RNA-seq, total RNA was isolated from nuclei preserved in DNA/RNA Shield and processed using a 3′-end counting RNA-seq protocol on an Illumina sequencing platform. Unique molecular identifiers (UMIs) were incorporated during library preparation to enable removal of PCR duplicates and accurate transcript quantification. Sequencing reads were aligned to the mouse reference transcriptome and collapsed by UMI to generate deduplicated gene-level count matrices. Differential gene expression analysis was performed using the processed count data provided by Plasmidsaurus.

Nucleosome-seq

For analyzing MNase-seq data, fastq files were processed with cutadapt -m 20 with the following adapter sequences: AGATCGGAAGAGCACACGTCTGAACTCCAGTCA, AGATCGGAAGAGCGTCGTGTAGGGAAAGAGTGT. Next, Bowtie2 was run with flags –very-sensitive, -X 1000, then converted to bam files with samtools, followed by samtools sort. Nucleosome occupancy was mapped with Danpos3 danpos.py dpos command to compare control versus CCl_4_-treated sample bam files with flags -m 1, -jd 120, -n F, -u 1e-10 to generate xlsx and wig files. The first 3 columns of the xlsx file were extracted to generated bed files. Then danpos.py profile command was run with wig file inputs and the bed positions extracted from dpos to generate genome-wide nucleosome occupancy maps with the flags: --genomic_sites center, --flank_up 500, --flank_dn 500. Chipseeker was used to annotate nucleosome positions in bed files with annotatePeak command (TxDb = TxDb.Mmusculus.UCSC.mm39.refGene, annoDb = org.Mm.eg.db, tssRegion = c(-500,500)). To generate nucleosome occupancy maps at differentially regulated genes we focused induced genes (log_2_FC > 1, q<0.05), repressed genes (log_2_FC<-1, q<0.05) and unchanged genes (q>0.8) as defined by bulk liver RNA-seq of control versus CCl_4_-treated mice. Unique gene symbols from these three groups were selected in the annotated dpos bed files and gene start or end positions were used as the TSS, depending on the strand. Nucleosome occupancy positions for nucleosomes within 300 bp of the TSS of differentially expressed genes were generated by running danpos.py profile command with flags: --genomic_sites center, --flank_up 500, --flank_dn 500.

rRNA-depletion RNA-seq

For analyzing data, fastq files were aligned with kallisto/0.46.0 using kallisto quant command and the mouse transcriptome index (mus_musculus.tar.gz/transcriptome.idx) from Pachter Lab (github.com/pachterlab/kallisto-transcriptome-indices/releases) with flag -b 100. Then sleuth was used to obtain differential expression from kallisto output files by running sleuth_prep, followed by sleuth_fit (so, ~group, ‘full’) to obtain the full model and sleuth_fit (so, ~1, ‘reduced’) to obtain the reduced model. Differential analysis testing with likelihood ratio test was performed with sleuth_lrt (so, ‘reduced’, ‘full’). Wald test results were obtained with sleuth_wt followed by sleuth_results to obtain comparisons. To generate signal tracks, fastq files were processed with STAR with flags –outSAMtype BAM SortedByCoordinate, --outSAMunmapped Within, --outSAMattributes Standard to generate bam files, which were indexed by samtools. BigWig files were created with deeptools bamCoverage command using bam as inputs. Graphics were generated with plotgardner library in R.

single-nuclei RNA-seq

CellRanger

The raw fastq files were processed using 10X CellRanger count command with mm39 reference genome and flags including –expect-cells=5000, --chemistry=SC3Pv4. This resulted in estimated nuclei counts as follows: control 4.8K, CCl_4_ 17.8K, CDKi 5.1K, CCl_4_+CDKi 3.5K.

CellBender

The CellRanger output files were used to run CellBender remove-background command with the following flags per sample: Control --expected-cells 5000, --total-droplets-included 15000, --cuda; CCl_4_ –expected-cells 10000, --total-droplets-included 30000, --cuda, --epochs 300, --learning-rate 0.000025; CDKi --expected-cells 2000, --total-droplets-included 15000; --cuda, --learning-rate 0.000025; CCl_4_+CDKi --expected-cells 1700, --total-droplets-included 13000, --cuda, --learning-rate 0.00005.

Remove Doublets

CellBender output was coerced into a Seurat object using SingleCellExperiment command and input to scDblFinder to remove doublets. This identified doublet rates of 7% in Control, 14.8% in CCl_4_, 6.5% in CDKi, 5.5% in CCl_4_+CDKi.

Cell Type Annotation

Annotation was performed with Liver Cell Atlas (livercellatlas.org) mouse StSt (All Liver cells) using the Seurat Read10X command with files: matrix.mtx.gz, barcodes.tsv.gz, features.tsv.gz, followed by CreateSeurateObject(counts) command. Then metadata was added from the LiverCellAtlas annot_mouseStStAll.csv file. Top expressed genes in each cluster were extracted from StSt using NormalizeData, then SetIdent, then FindAllMarkers with flags only.pos=TRUE, min.pct=0.25, logfcthreshold=0.25. The output top expressed genes were clustered by group, ordered by log2FC and used as the marker_list for annotation. Annotation of our data was performed with Seurat AddModuleScore for each sample with flags: --features=marker_list, nbin=10, ctrl=25. The meta.data of each Seurat was renamed from cluster numbers to cell type annotation. Next, a series of Seurat commands were run to normalize, scale and cluster the data. These included: NormalizeData, FindVariableFeatures, ScaleData, RunPCA, RunUMAP, FindNeighbors all with default parameters and FindClusters with resolution=0.5. Cell clusters were then assigned based on their most prominent cell type, which was calculated by average marker gene expression minus the average control gene expression using the expression summarise(across(all_of(celltype_cols), mean)) from dply.

Integration of datasets

All four datasets were merged with merge command, then the cell barcodes were renamed to ensure uniqueness. Then data were re-normalized, scaled and clustered with the following Seurat commands: NormalizeData, FindVariableFeatures, ScaleData, RunPCA with default parameters. Integration was acheived with Seurat IntegrateLayers command with method=CCAIntegration. Then clustering was performed with FindNeighbors, followed by Findclusters with resolution=0.4, then RunUMAP. Validation of the integration was confirmed by visualizing UMAP labeled by either cell type or condition, using the Seurat DimPlot command.

Differential Expression

Only hepatocyte cell types were used for differential expression, which were selected from the integrated dataset. The Seurat command FindMarkers was used to compare Control sample versus the other three samples with test.use=”wilcox”, logfc.threshold=0, min.pct=0.1. AnnotationDbi R package was used to annotate genes. Then enrichGO command from ClusterProfiler was run with ont=”BP”, pAdjustMethod=”BH”, pvalueCutoff=0.05, qvalueCutoff=0.05 on both upregulated and downregulated gene sets selected with logFC <-1 or >1 and qval 0.05. KEGG enrichment was performed with enrichKEGG from ClusterProfiler with universe=background_entrez, pvalueCufoff=0.05, qvalueCutoff=0.05 on similarly selected gene sets.

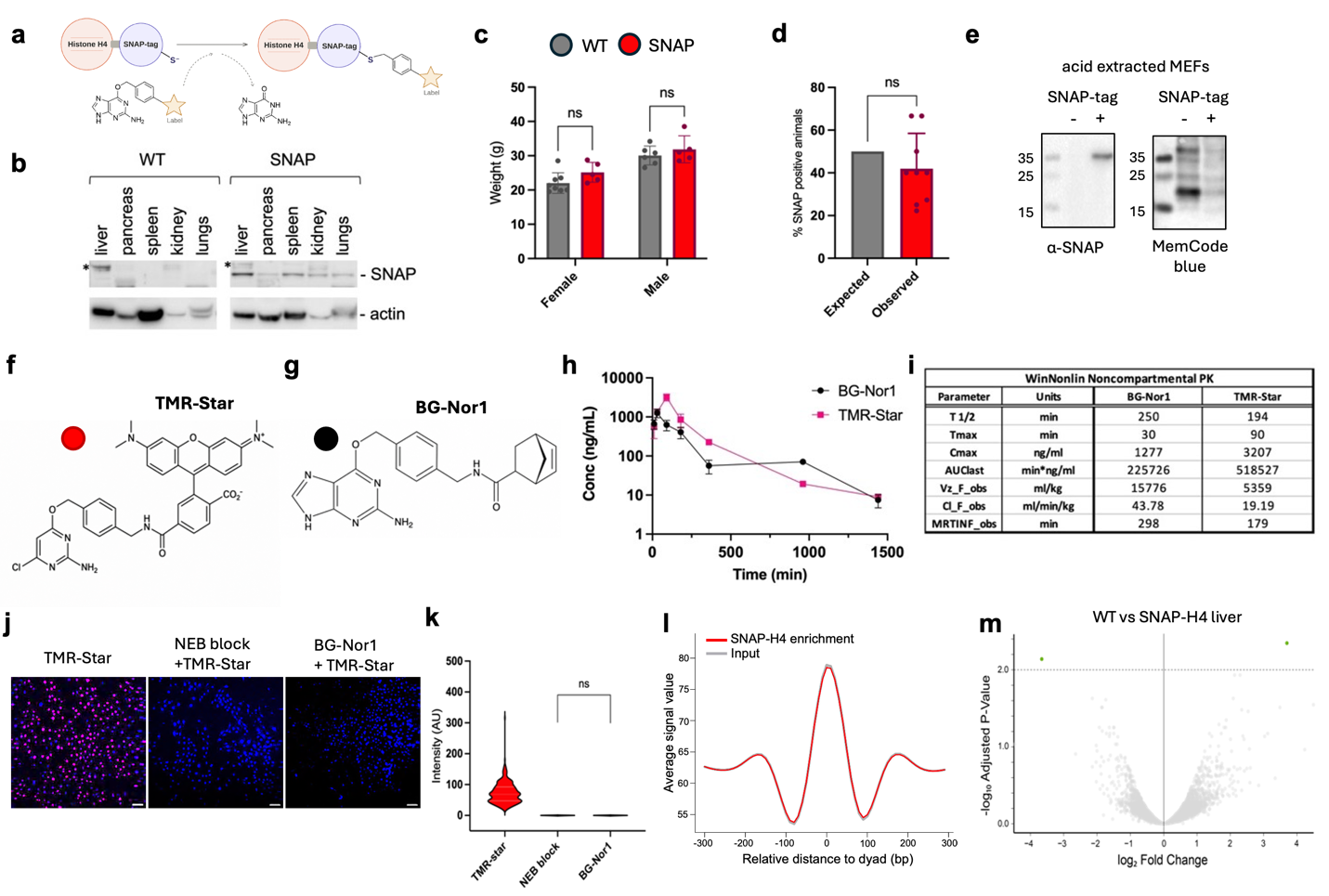

Fig. S1. a, Schematic of SNAP tag covalent adducts formed from benzylguanine analogs. b, Immunoblot confirmation of SNAP–H4 protein expression across multiple tissue; n = 1, *nonspecific. c, Body weights of 10-week-old littermates (WT vs. SNAP-H4); n = 8. d, Frequency of SNAP–H4 positive offspring compared with expected Mendelian inheritance from 9 breeder pairs. e, Mouse embryonic fibroblasts (MEFs) acid-extracted to validate presence of SNAP-tag integrated into nucleus by immunoblot. f & g, Chemical structures of SNAP substrates TMR-star & BG-Nor1 synthesized at scale for *in vivo* use. h & i, Pharmacokinetics studies (i.p.) of TMR-star and BG-Nor1 analyzed using Phoenix WinNonlin. j, labeling of histones *in vitro* with TMR-star on MEFs derived from *Snap-H4c3* animals, along with validation of BG-Nor1 and k, their quantification; n = 2, scale bar 10 μm, imaged at 20x magnification. l, *Snap-H4c3* MEFs subjected to MNase digestion & SNAP-BG bead pulldown nucleosome seq (n = 2), input vs SNAP-H4 enrichment shows global SNAP-H4 nucleosome occupancy. m, RNA-seq differential gene expression of WT vs *Snap-H4c3* liver tissue (n = 3).

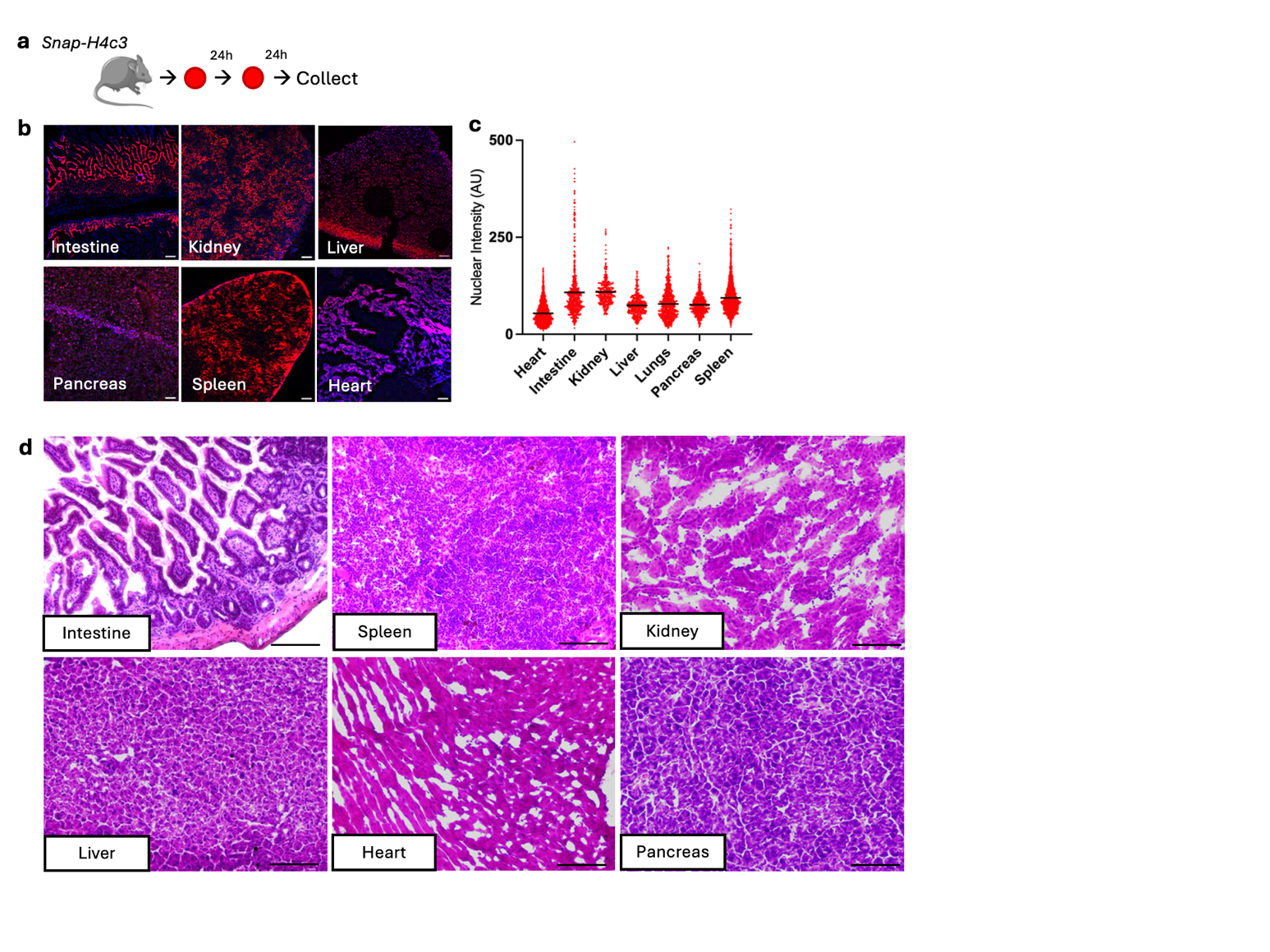

**Fig. S2.** **a**, Experimental schematic for measuring SNAP-H4 labeling with TMR-star serial injections *in vivo*. *Snap-H4c3* mice received TMR-star followed by TMR-Star 24 h later. Tissues were harvested 24 h later, flash frozen, cryo-sectioned, stained with DAPI mounted & imaged for 570 nm fluorescence by confocal. **b**, Representative fluorescence images of indicated tissues showing tissue-specific SNAP-H4 labeling with DAPI nuclear counterstain; n =2 biological, n = 3 technical; scale bar 10 μm, imaged at 10x magnification**. c**, Quantification of nuclear TMR fluorescence intensity across indicated tissues (n = 2). Each point represents an individual nucleus. **d**, Representative hematoxylin and eosin (H&E)-stained tissue sections from the indicated experimental groups to evaluate tissue morphology. Images shown are representative of the analyzed samples from Fig.1e,f; scale bar 10 μm, imaged at 20x magnification.

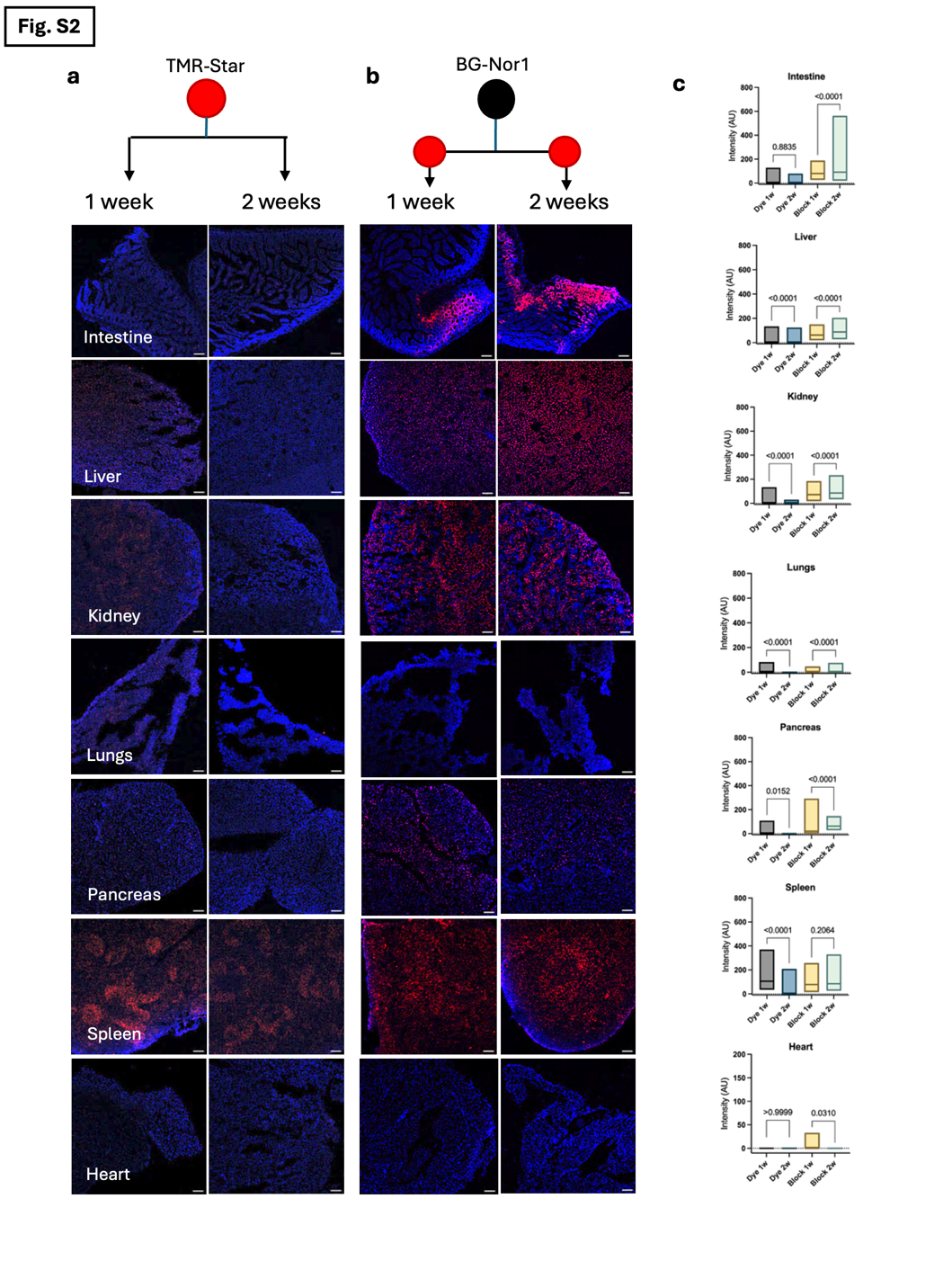

**Fig. S3**. **a**, Experimental schematic for assessing the long-term residence of TMR-Star–labeled histones followed by the results. *Snap-H4c3* mice received a single TMR-Star (5 mg/kg) injection and tissues were harvested either 1 or 2 weeks later, flash frozen, cryo-sectioned, stained with DAPI mount & imaged for 570 nm fluorescence confocal; n = 2 biological replicates, scale bar 10 μm, imaged at 20x magnification **b**, Experimental schematic for assessing the duration of BG-Nor1 blocking activity *in vivo*. Mice were administered BG-Nor1, followed by a 1- or 2-week waiting period before TMR-Star administration 24 h prior to tissue collection. Tissues were flash frozen, cryo-sectioned, stained with DAPI mount & imaged for 570nm fluorescence confocal; n = 2 biological replicates, scale bar 10 μm, imaged at 20x magnification. **c**, Tissue-wise quantification of fluorescence intensity across all analyzed organs; n = 2, 2-way ANOVA, Šídák's multiple comparisons test.

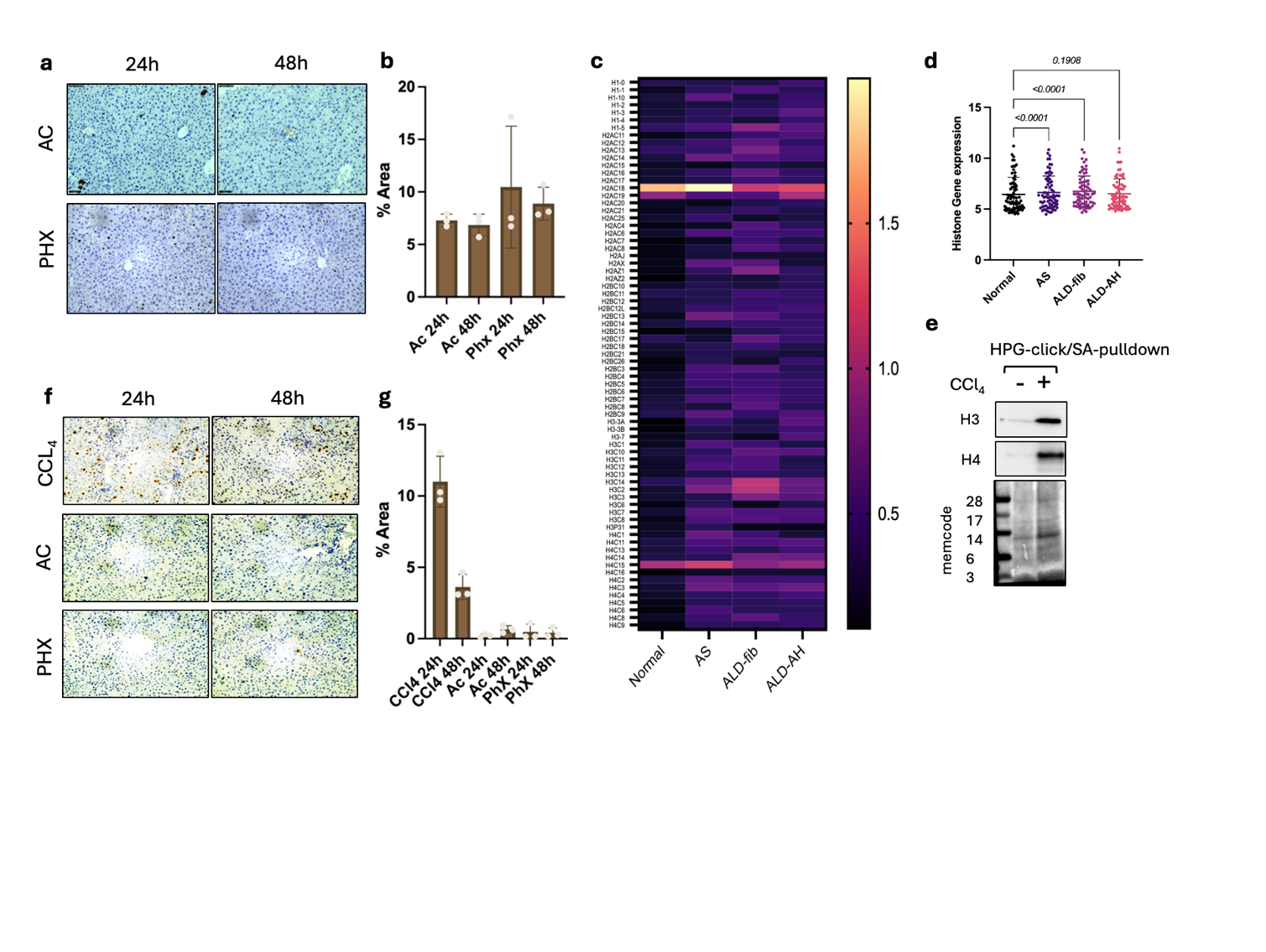

**Fig. S4.** **a**, Representative Ki67 immunohistochemistry (IHC) of liver sections collected 24 hours and 48 hours following acetaminophen (Ac) or partial hepatectomy (PHX); scale bar 10 μm, imaged at 20x magnification. **b**, Quantification: DAB-positive Ki67 staining was quantified as the percentage of positively stained area relative to the total tissue area (area %). **c**, Heatmap of differentially expressed histone mRNA transcripts from human livers; normal, alcohol-related steatosis (AS), alcoholic liver disease (ALD) with fibrosis (ALD-fib), and ALD with alcoholic hepatitis (AH) . **d**, quantification of gene expression from (c); Welch’s t-test, n = 10 (normal), 10 (AS), 13 (ALD-fib), 28 (ALD-AH). **e**, HPG-biotin-click assay on CCl_4_ treated and untreated liver extracts; streptavidin-pull down, SDS-PAGE, transferred to nitrocellulose membrane and detected with streptavidin-HRP chemiluminescence. **f**, gH2AX IHC of liver sections following acute CCl₄ injury, AC, or PHX at 24 hours and 48 hours; scale bar 10 μm, imaged at 20x magnification. **g**, Quantification: DAB-positive gH2AX staining was quantified as the percentage of positively stained area relative to the total tissue area (area %).

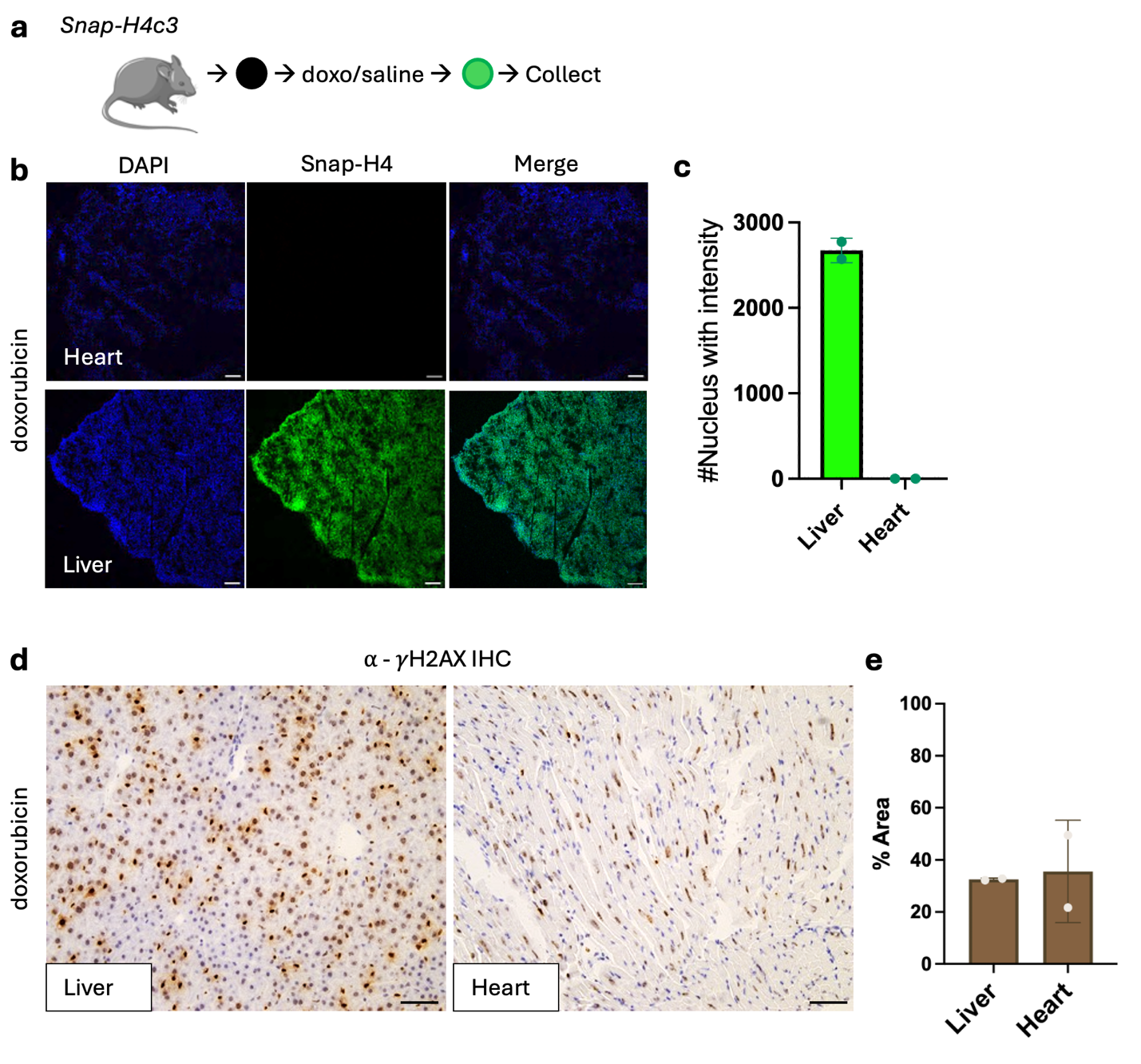

**Fig. S5. a**, Experimental schematic showing doxorubicin treatment design. *Snap-H4c3* mice were first blocked with BG-Nor1, treated with doxorubicin (20 mg/kg) or saline control, labeled with BG-Oregon (5 mg/kg) green, and subsequently harvested. **b**, Representative liver sections showing SNAP-H4 labeling following doxorubicin treatment at 24 hours in heart and liver tissues; scale bar 10 μm, imaged at 10x magnification. **c**, BG-Oregon green positive nuclei with intensity were quantified, n = 2, one-way ANOVA Brown-Forsythe test. **d**, Representative gH2AX immunohistochemistry (IHC) of heart and liver sections collected 24 h following doxorubicin treatment; scale bar 10 μm, imaged at 20x magnification **e**, quantification n = 2 biological replicates; DAB-positive gH2AX staining was quantified as the percentage of positively stained area relative to the total tissue area (area %).

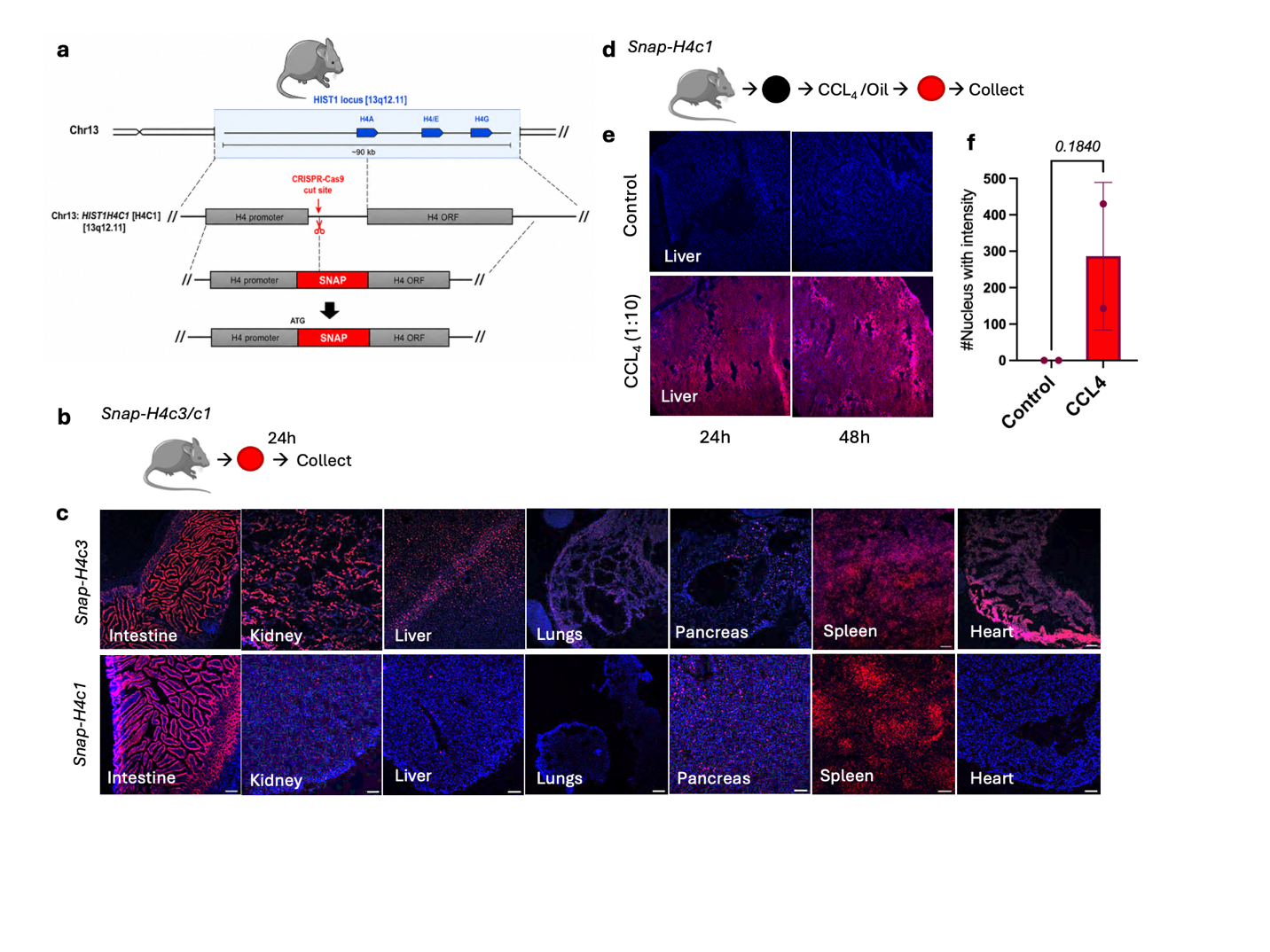

**Fig. S6. a,** Schematic of SNAP knock-in strategy into *H4C1* locus. **b**, Experimental schematic showing TMR-star administration to *Snap-H4c1* versus *Snap-H4c3* mice, tissues harvested after 24 h. **c**, Representative tissue sections of SNAP-H4 labeling shows tissue expression based on which histone gene was tagged. Tissues were harvested, flash frozen, cryo-sectioned, stained with DAPI & imaged at 570 nm by confocal; n = 2, scale bar 10 μm, imaged at 10x magnification **d**, *Snap-H4c1* mice were first blocked with BG-Nor1, treated with CCl_4_ or oil control, labeled with TMR-star, and subsequently harvested. **e**, Representative liver sections showing SNAP-H4 labeling following CCl_4_ treatment. Tissues were harvested, flash frozen, cryo-sectioned, stained with DAPI & imaged at 570 nm by confocal **f**, Quantification of labeling in *Snap-H4c1* mice post CCl_4_ treatment; TMR-positive number of nuclei with intensity were quantified, n = 2, one-way ANOVA Brown-Forsythe test.

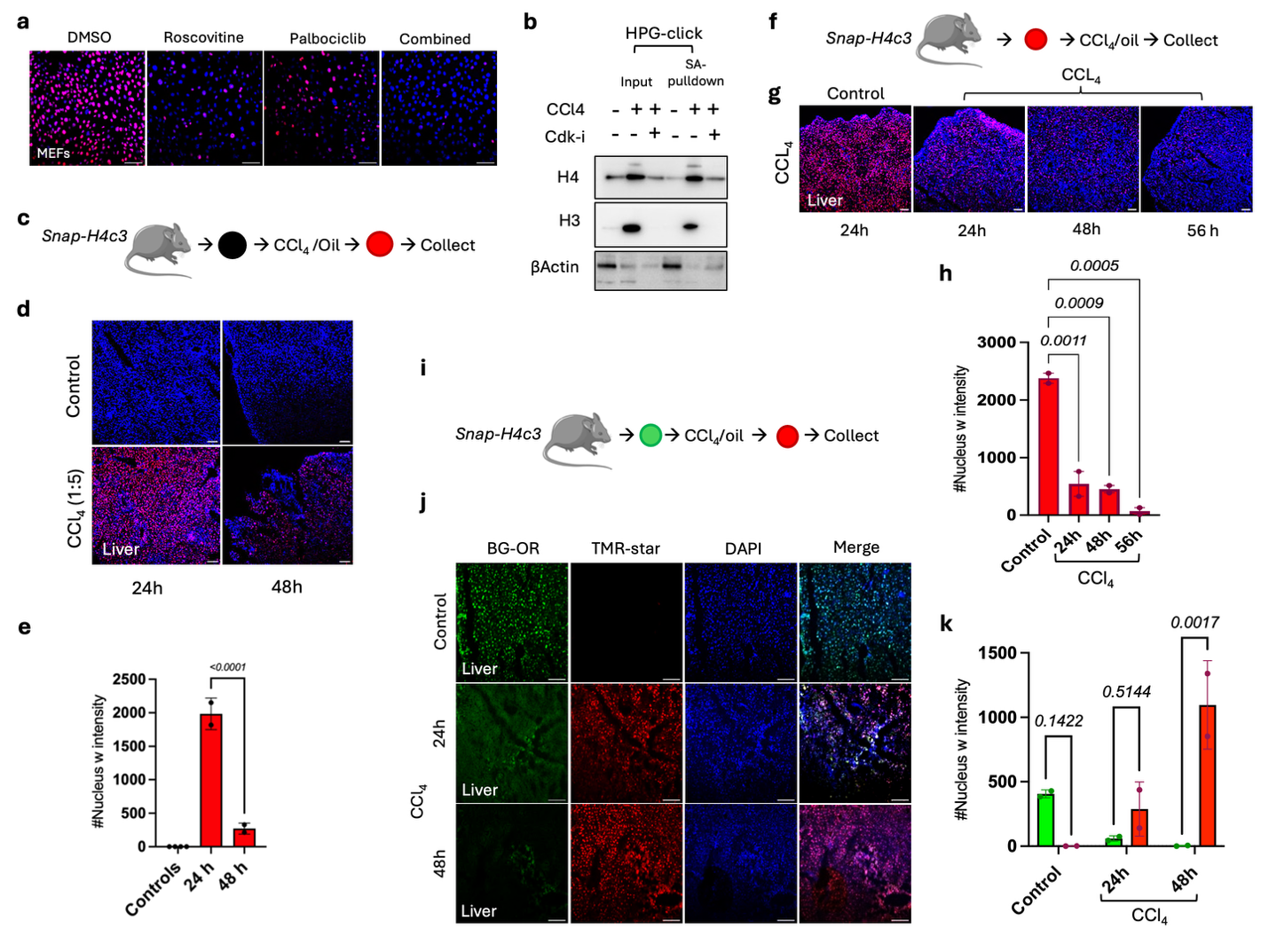
 **Fig. S7. a**, *Snap-H4c3* MEFs treated with 20 μM CDK-inhibitors. Cells were plated on coverslips, blocked with BG-Νor1 1 hour before treating with CDK inhibitors (palbociclib, roscovitine). After 24 hours, cells were labeled with TMR-star, fixed, permeabilized, DAPI stained and imaged by ZEISS confocal (n = 3), scale bar 10 μm, 20x magnification. **b**, HPG-biotin-click assay on CCl_4_ only and CCl_4_ + CDK-i combo treated liver extracts; streptavidin-pull down, SDS-PAGE, transferred to nitrocellulose membrane, probed for total histone H4, H3 and b-actin. **c**, Experimental schematic showing SNAP-H4 labeling at higher CCl_4_ dose (1:5). *Snap-H4c3* mice were first blocked with BG-Nor1, treated with CCl₄ or oil control, labeled with TMR-Star, and subsequently harvested. **d**, Representative liver sections showing SNAP-H4 labeling following CCl₄ treatment at 24 hours and 48 hours; scale bar 10 μm, imaged at 10x magnification. **e**, TMR-positive number of nuclei with intensity were quantified, n = 2, one-way ANOVA Brown-Forsythe test. **f & g**, Schematic and results of treating *Snap-H4c3* mice first with TMR-star followed by CCl_4_(1:10); tissues were harvested, flash frozen, cryo-sectioned, stained with DAPI & imaged at 570 nm by confocal; scale bar 10 μm, imaged at 10x magnification. **h**, TMR-positive number of nuclei with intensity were quantified; n =2, ordinary one-way ANOVA. **i & j**, Dual color CCl_4_ experimental set-up and results; *Snap-H4c3* mice were first labeled with BG-Oregon green followed by CCl_4_ treatment for 24 & 48 hours, followed by TMR-star labeling; tissues were harvested, flash frozen, cryo-sectioned, stained with DAPI & imaged at 514 nm and 570 nm by confocal; scale bar 10 μm, 10x magnification. **k**, TMR and Oregon green positive number of nuclei with intensity were quantified; n = 2, 2-way ANOVA.
